# The blackgrass genome reveals patterns of divergent evolution of non-target site resistance to herbicides

**DOI:** 10.1101/2021.12.14.472569

**Authors:** Lichun Cai, David Comont, Dana MacGregor, Claudia Lowe, Roland Beffa, Christopher Saski, Paul Neve

## Abstract

Globally, weedy plants result in more crop yield loss than plant pathogens and insect pests combined. Much of the success of weeds rests with their ability to rapidly adapt in the face of human-mediated environmental management and change. The evolution of resistance to herbicides is an emblematic example of this rapid adaptation. Here, we focus on *Alopecurus myosuroides* (blackgrass), the most impactful agricultural weed in Europe. To gain insights into the evolutionary history and genomic mechanisms underlying adaptation in blackgrass, we assembled and annotated its large, complex genome. We show that non-target site herbicide resistance is oligogenic and likely evolves from standing genetic variation. We present evidence for divergent selection of resistance at the level of the genome in wild, evolved populations, though at the transcriptional level, resistance mechanisms are underpinned by similar patterns of up-regulation of stress- and defence-responsive gene families. These gene families are expanded in the blackgrass genome, suggesting that the large, duplicated, and dynamic genome plays a role in enabling rapid adaptation in blackgrass. These observations have wide significance for understanding rapid plant adaptation in novel stressful environments.

## Main

Human-mediated environmental change is driving rapid evolutionary responses in the global biota ^1,2^ and it is important to understand the outcome of these changes in natural and agricultural plant populations and communities. Plant genomes offer glimpses into the adaptive potential of plant populations when challenged with novel environmental stresses. Agricultural weeds rapidly adapt in managed agroecosystems and have been proposed as models to address fundamental questions in plant ecology and evolution ^3-7^. Their global impacts on crop yields provides an additional economic incentive to understand weed adaptation.

Herbicide use has become a mainstay of weed management in most industrialized agricultural economies. Unsurprisingly, heavy reliance on herbicides has resulted in the rapid and widespread evolution of resistance, making herbicide resistance a widely studied weedy trait ^8-9^. Two main ‘types’ of herbicide resistance are recognized ^10-11^. Target site resistance (TSR) refers to modification of the sequence, copy number or expression of the gene encoding the herbicide target enzyme. Non-target site resistance (NTSR) encompasses a range of mechanisms that limit herbicide delivery to its site of action. Typically, NTSR is inherited in a quantitative manner, and despite some advances in identifying and/or validating causal loci ^12-15^, efforts to discern the genomic basis and evolutionary dynamics of this trait have been hampered by lack of access to genomic resources in target species.

The diploid, allogamous grass, *Alopecurus myosuroides* (blackgrass) is native to the Eastern Mediterranean and West Asia ^16^. It is now a widespread and impactful weed in agricultural crops in the UK ^17^, France ^18^ and Germany ^19^, with evidence of an ongoing range expansion in Europe. *Alopecurus* species are also major weeds in China ^20^. Blackgrass populations appear to be uniquely prone to the rapid and widespread evolution of herbicide resistance. In a nationwide survey in England conducted, most blackgrass populations exhibited resistance to multiple herbicide modes of action ^21^. Resistance was conferred by both TSR and NTSR mechanisms that often co-existed in populations, and there was evidence that current and historical herbicide use regimes favoured the evolution of the generalist NTSR mechanism ^22^. Herbicide resistant blackgrass is estimated to cost UK farmers £0.4 billion per year ^23^.

Access to genomes and genomic resources for weed species will greatly enhance the capacity to unravel pattern and process in economically and ecologically important weedy plant species ^24^. Here, we present a high-quality reference genome of blackgrass. We analyse genome structure and function to infer genomic features that may predispose blackgrass to the rapid evolution of weediness and present data from genomic and transcriptomic resequencing of two well-characterised NTSR populations.

## Results

### Genome assembly and annotation

Genome analysis indicated that blackgrass (*A. myosuroides*) has a large genome (3.31-3.55 Gb) and exhibits heterozygosity of 1.52% and repeat content of 84.2% contributing to the large genome size (Supplementary Tables 1 and 2). We adopted a hierarchical sequencing approach that includes complementary single-molecule sequencing/mapping technologies coupled with deep coverage short read sequences to generate a pseudo-chromosome reference genome assembly for blackgrass (Supplementary Figure 1). The total primary contig length is 3,475 Mb, which is consistent with our genome size estimations based on flow cytometry and k-mer analysis (3,312-3,423 Mb and 3,400-3,550 Mb, respectively). The final polished blackgrass genome assembly size was highly contiguous at 3,572 Mb, including 3,400 (95.2%) Mb ordered as seven pseudo-chromosomes with only 172 Mb of unanchored sequences (Table1, Supplementary Table 3).

**Table 1.**
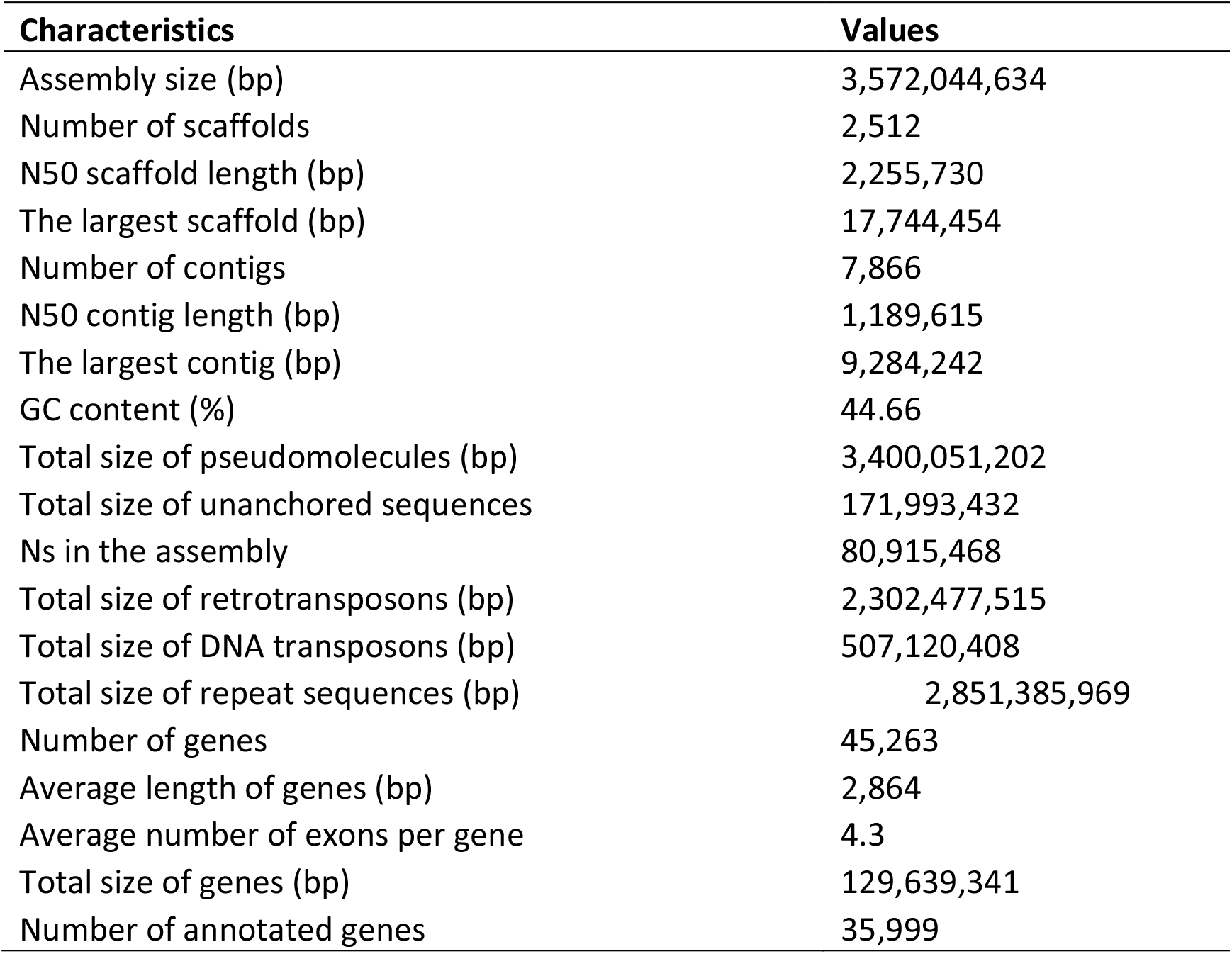
Assembly statistics of the blackgrass genome.

| Characteristics | Values |
| --- | --- |
| Assembly size (bp) | 3,572,044,634 |
| Number of scaffolds | 2,512 |
| N50 scaffold length (bp) | 2,255,730 |
| The largest scaffold (bp) | 17,744,454 |
| Number of contigs | 7,866 |
| N50 contig length (bp) | 1,189,615 |
| The largest contig (bp) | 9,284,242 |
| GC content (%) | 44.66 |
| Total size of pseudomolecules (bp) | 3,400,051,202 |
| Total size of unanchored sequences | 171,993,432 |
| Ns in the assembly | 80,915,468 |
| Total size of retrotransposons (bp) | 2,302,477,515 |
| Total size of DNA transposons (bp) | 507,120,408 |
| Total size of repeat sequences (bp) | 2,851,385,969 |
| Number of genes | 45,263 |
| Average length of genes (bp) | 2,864 |
| Average number of exons per gene | 4.3 |
| Total size of genes (bp) | 129,639,341 |
| Number of annotated genes | 35,999 |

Both the euchromatic and heterochromatic components of the blackgrass genome are highly complete as supported by BUSCO scores (96.9% from the *Embryophyta* lineage) ^25^ and a high long terminal repeat assembly index across the genome (LAI - 9.6-35.2) ^26^, with a mean value of 21.9 (Supplementary Table 4; Supplementary Figure 4). In addition, the Illumina short reads (81×) returned a 99.6% mapping rate and covered 99.9% of the assembled genome. We identified 8,026,403 polymorphisms as SNPs or InDels (Figure 1a), which was expected from the predicted heterozygosity level of the blackgrass genome. We also observed a high correlation (r = 0.98) between the assembled chromosome and cytogenic chromosome lengths based on published data ^27^.

**Figure 1.**
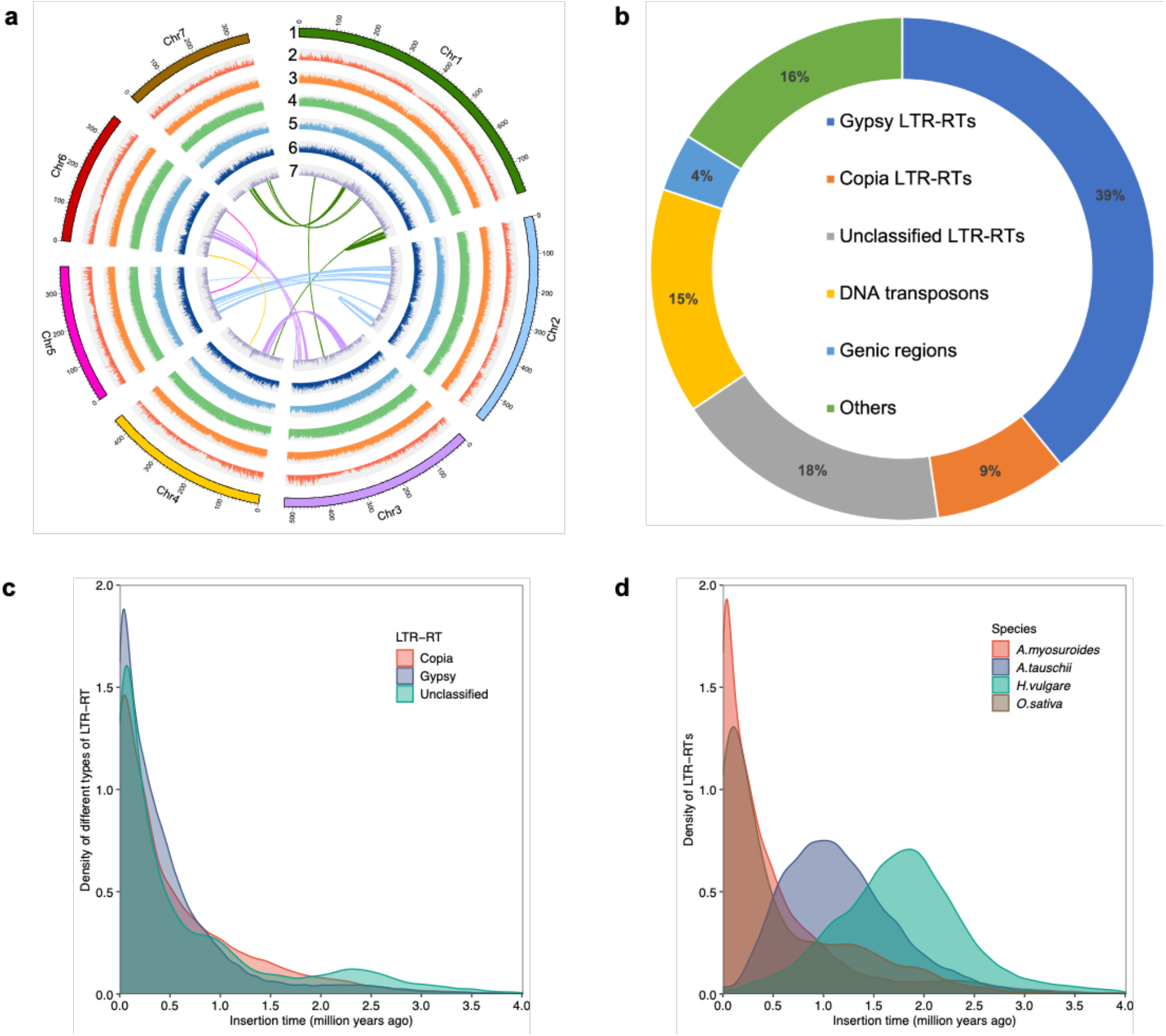
Genomic features and components of the *A. myosuroides* genome. **a**, overview of the *A. myosuroides* genome, including the assembled seven chromosomes (1), distribution of protein-coding genes (2), distribution of GC content across the genome (3), distribution of transposable elements (4), distribution of *Gypsy* family of long terminal repeats retrotransposons (5), distribution of *Copia* family of long terminal repeats retrotransposons (6), distribution of SNP/Indel (7). All the histograms (from 1 to 7) were featured in a 1-Mb sliding window. Connecting line in the center of the diagram represents a genomic syntenic region covering at least 10 paralogues. **b**, Proportions of the major elements in the blackgrass genome, including *Gypsy* LTR-RTs, *Copia* LTR-RTs, unclassified LTR-RTs, DNA transposons, coding DNA and unannotated sequences. **c**, The insertion time distribution of different types of LTR-RT in the blackgrass genome. **d**, The insertion time distribution of intact LTR-RTs in the blackgrass genome compared to those in goatgrass (progenitor of the wheat D genome), barley and rice.

We annotated 45,263 protein-coding genes based on *de novo* and homology-based predictions and transcriptome data from multiple tissues (Supplementary Figure 3). Mean gene length was 2,864 bp, with an uneven distribution across the chromosomes with increased gene density toward the distal ends that recedes to very low density in the middles of chromosomes (Figure 1a). Among these protein-coding genes, 2,385 were annotated as transcription factors. In addition, 4,258 non-coding RNAs were identified, including 1,369 transfer RNAs, 941 ribosomal RNAs, 513 micro RNAs and 1,425 small nuclear RNAs (Figure 1a for genome overview).

### Transposon elements and the burst of LTR retrotransposons

We annotated 2,851 Mb (81.7%) of sequence in the assembled genome as transposable elements (TEs) (Supplementary Table 5). A total of 5,287,231 repeat elements were identified and the dominant type of TE was long terminal repeat retrotransposons (LTR-RTs), representing approximately 80.3% (2290 Mb) of annotated TEs and amounting to 65.6% of the blackgrass genome size. *Gypsy, Copia* and unclassified retrotransposon elements contributed to 39.2%, 8.6% and 17.9% of the genome size, respectively. DNA transposons contributed to 14.5% of the genome and the CACTA transposon were the most abundant DNA transposons, accounting for 5.9% of the annotated TEs and 4.9% of the assembled blackgrass genome (Figure 1b).

Retrotransposons are highly unstable and have played an important role in the evolution of plant genomes ^28^. We observed a single peak of insertion time, occurring approximately 0.1 million years ago (Ma), for *Gypsy, Copia*, and unclassified retrotransposons in blackgrass, which suggests a recent burst of LTR retrotransposons in the genome (Figure 1c). In addition, we observed a higher proportion of recent LTR-RT insertions when compared to those in rice, barley and goatgrass (progenitor of the wheat D genome). Moreover, the burst of retrotransposons in blackgrass was more recent than those in barley (*Hordeum vulgare*) and goatgrass (*Aegilops tauschii*) but occurred at a similar time in blackgrass (*A. myosuroides*) and rice (*Oryza sativa*) (Figure 1d). Therefore, the recent large-scale burst of retrotransposons might have contributed to blackgrass genome expansion, explaining the large genome size.

### Phylogenetics and gene expansion and contraction

To assess the divergence time between blackgrass and other grasses, we constructed a phylogenetic tree based on the concatenated sequence alignment of the 476 single-copy orthologous genes shared by blackgrass and 11 other species (Figure 2a). The divergence between blackgrass and barley was after the separation of blackgrass from rice and *Brachypodium distachyon*. The divergence time between blackgrass and barley was estimated at 37.9 million years ago.

**Figure 2.**
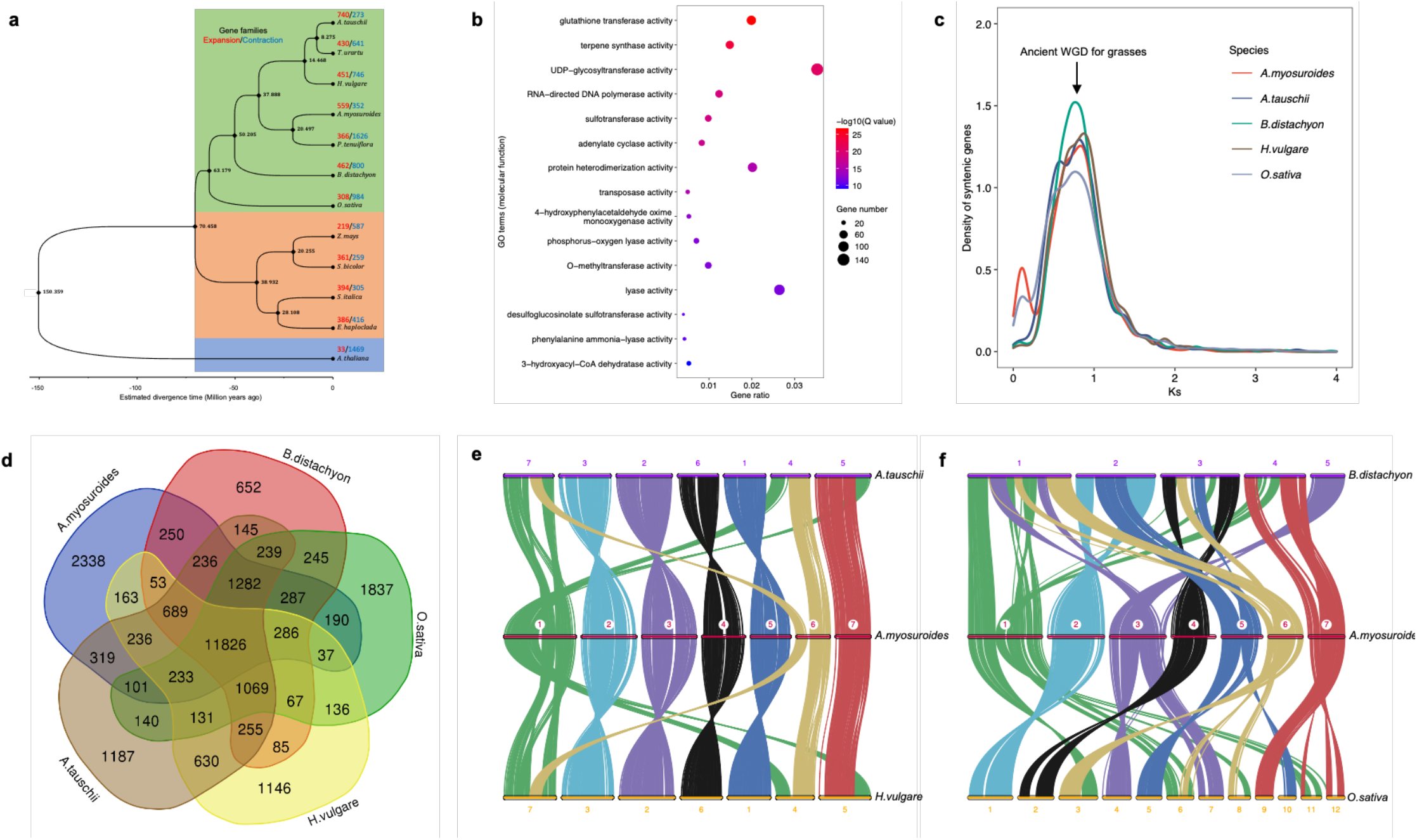
Evolution and Comparative genomics of the *A. myosuroides* genome. **a**, Phylogenetic tree of 12 plant species and gene family expansion and contraction. Inferred divergence time is denoted at each node in black. The red and blue numbers above the species name indicate the total number of expanded and contracted gene families, respectively. **b**, Gene Ontology (GO) enrichment analysis of expanded gene families in the blackgrass genome (molecular function category). **c**, The frequency distribution of synonymous substitution rates (Ks) of paralogous genes within each genome. A shared whole genome duplication event for grasses was assigned to the peak. **d**, Venn diagram of shared and unique gene families among five closely related Poaceae species. Each number represents the number of gene families. **e**, Syntenic blocks between blackgrass and other sequenced grass genomes, including goatgrass and barley. **f**, Syntenic blocks between blackgrass and other sequenced grass genomes, including *Brachypodium* and rice.

We also examined gene family evolution through expansion and contraction events. A total of 33,757 orthologous gene families composed of 382,550 genes were identified from 12 species, of which 6,470 gene families were shared by all the species. Blackgrass contains 1,678 species-specific gene families, which is the most among all the investigated species (Supplementary Figure 4). A total of 559 and 352 gene families were identified with significant expansion and contraction, respectively. GO enrichment analysis of the expanded genes revealed that they were mainly related to multiple enzymatic functions, including glutathione transferase (GST), UDP-glycosyltransferase (UGT), and monooxygenases, all of which have been reported to be associated with non-target site herbicide resistance (Figure 2b).

### Genome duplication and comparative genomics

We explored evidence for whole genome duplication events in the blackgrass genome. Synonymous substitution rate (*K*_*s*_) values were calculated from 1,884 paralogous gene pairs and were used to infer the age distribution of the duplication events, which are evident with two distinct peaks at *K*_*s*_ values of 0.1 and 0.8, respectively (Figure 2c). To determine if these peaks were species-specific or common in the grass family, we performed the same analysis for rice, barley, *Brachypodium* and goatgrass (Figure 2c). The results indicated that the peak at 0.8 was shared in all investigated species, suggesting blackgrass underwent the same ancient whole genome duplication in the ancestor of *Poaceae* species ∼70 MYA ^29^. The peak at 0.1 is not apparent in other species, suggesting that this genome duplication event is unique for blackgrass. We further examined the paralogous genes within the duplication events and found that the peak at 0.1 was evidenced by a high density of ‘co-located’ paralogous genes on chromosomes 1, 2, and 3 (Figure 1a) which suggests the blackgrass genome underwent some small-scale local duplication events in addition to the whole genome duplication. To investigate gene duplication structure in the blackgrass genome, we analysed all chromosome-anchored protein-coding genes for duplications and organization. The results show that blackgrass contains 9,106 singletons, 20,856 dispersed duplicated genes, 4,607 proximally duplicated genes, 4,967 tandemly duplicated genes and 3,615 segmentally duplicated genes.

Conserved genome structure and organization between blackgrass, barley, goatgrass, *Brachypodium*, and rice was examined through collinearity and macro-/micro-synteny approaches. We identified 11,826 plant gene families shared by all five species with 2,338 blackgrass-specific gene families (Figure 2d). Blackgrass chromosomes 2, 3, 4, 5 and 7 are completely collinear with barley chromosomes 3, 2, 6, 1, and 5, respectively (Figure 2e). For blackgrass chromosome 1, most regions were colinear with barely chromosome 7, with two small regions being colinear with parts of barley chromosome 4 and 5. Blackgrass chromosome 6 was colinear with barely chromosome 4 and a small part of chromosome 1 indicating that blackgrass genome content and structure most closely resembles that of barley. We observed a similar pattern of collinearity between blackgrass and goatgrass as we did between blackgrass and barley (Figure 2e). To investigate the chromosome evolution of blackgrass, we used rice as the reference species for comparison because it has retained 12 ancestral grass karyotype chromosomes. We found that blackgrass chromosomes 2 and 4 were derived from single ancient chromosomes 1 and 2, respectively (Figure 2f). All other blackgrass chromosomes were derived from fusion events between ancient chromosomes, including the large blackgrass chromosome 1 derived from fusion events among ancient chromosomes 3, 6, 8, 11 and 12; 3 from 4 and 7; 5 from 5 and 10, 6 from 3, 6 and 8; 7 from 9, 11 and 12. These chromosome reshuffling events might have contributed to the introduction of genetic variation and speciation of blackgrass. Combined, these results indicate that blackgrass diverged from *Brachypodium* and rice earlier than barely and goatgrass.

### QTL-seq bulk segregant analysis for NTSR

To identify the genomic regions controlling herbicide resistance, we performed bulk segregant analysis in the CC2 and CC5 families to identify ΔSNP values with trait significance ^30,31^. We obtained 3,402,057 and 3,205,888 reliable SNPs for each of the CC2 and CC5 families, respectively (Supplementary sFigure 5). We identified 7 significant QTLs in the CC2 family distributed among chromosomes 2,3,5, and 6 (Supplementary Table 6). In the CC5 family we identified 8 QTLs distributed mainly on chromosome 3 with 1 region on chromosome 2 (Supplementary Table 6). Interestingly, there was no overlap between QTL regions identified in the two seed families, however 12 of the 15 total QTL regions were located on chromosomes 2 and 3 (Figure 3). These two chromosomes also showed the greatest density of differentially expressed genes (DEGs), with almost half (33) of the 68 consistent DEGs located on these two chromosomes, along with half of the previously reported NTSR candidate loci for this species (Figure 3). These results suggest that chromosomes 2 and 3 are ‘hot-spots’ for NTSR evolution in this species. In addition, a total of 371 genes were encoded within the 15 identified QTLs, with each QTL containing between 10 and 58 genes. Among the 15 identified QTL regions, seven contain differentially expressed genes identified between susceptible and resistant plants; six of them contain transcription factors. The most significant QTL was identified on chromosome 2 in the CC2 family, which covered 2.5 Mb and contains 33 candidate genes. An NADPH-dependent aldo-keto reductase gene was present in this region and was upregulated in resistant plants for both CC2 and CC5 families. Members of this gene family have been reported to be associated with non-target herbicide resistance in other weed species ^32^.

**Figure 3.**
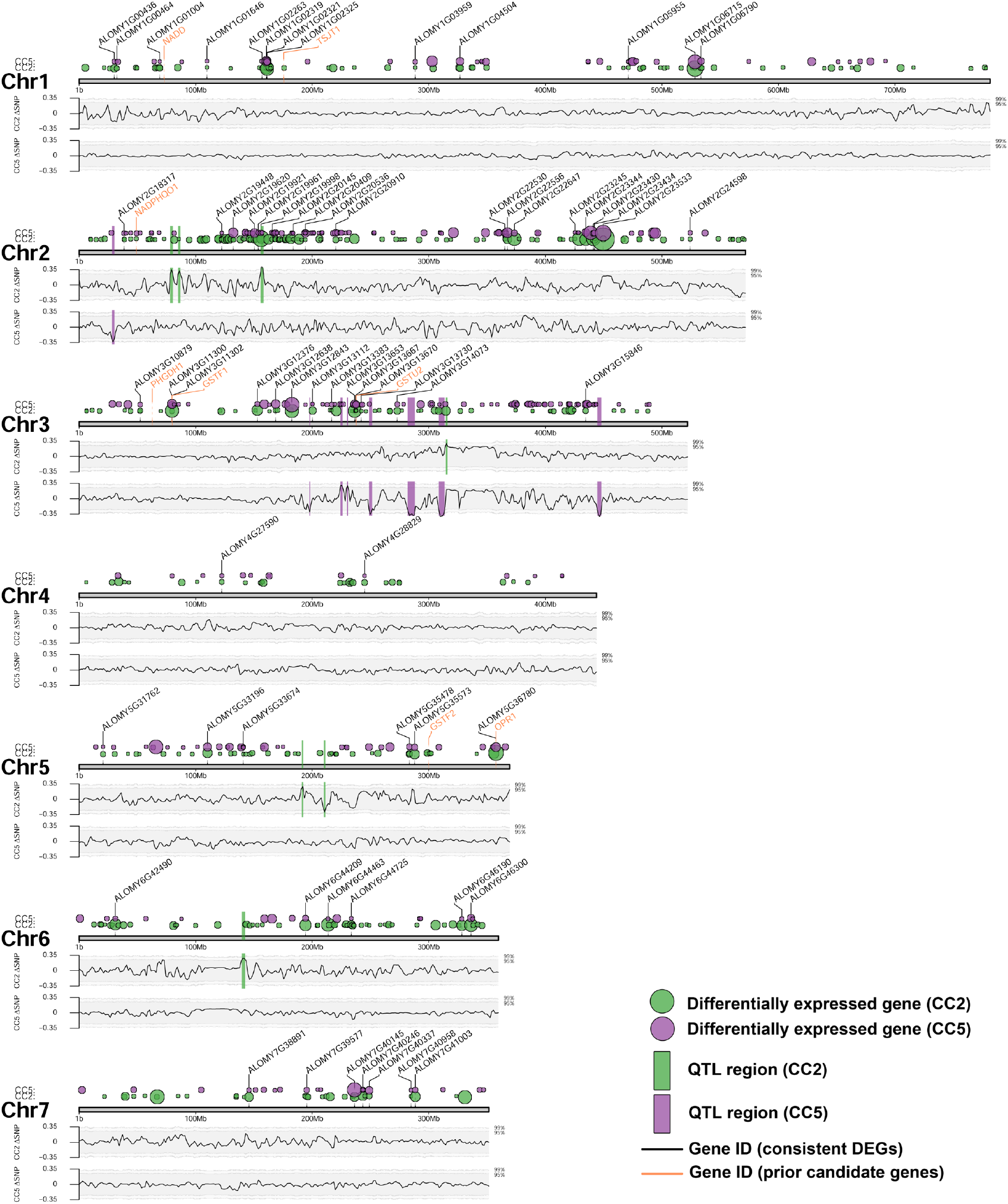
Location across the genome of the differentially expressed genes associated with the NTSR trait. Green and purple circles show the position of DEGs identified in the CC5 and CC2 seed families respectively. Circle sizes are relative to the adjusted P-value, whereby larger circles denote stronger significance. DEGs consistent across both families are marked with black labels, while orange labels show the position of previously reported NTSR candidate genes. Lower sections show the change in ΔSNP index across these chromosomes for the CC2 (top) and CC5 (bottom) families. Shaded regions represent the 95% and 99% confidence bounds for each SNP. Vertical green and purple bars show the QTL regions for the CC5 and CC2 families, identified from their ΔSNP index.

### RNA-seq analysis of herbicide resistance

To identify deferentially expressed genes between susceptible and resistant plants, we performed RNA-seq analysis in two seed families (CC2 and CC5). Principal components analysis of gene expression data (19,937 genes across 19 biological samples) indicates both seed families and resistance phenotypes contain significant sources of variation between samples (Figure 4a). Seed family (CC2 vs. CC5) was the stronger source of variance accounting for ∼58% of the total variance on the first Principal Component (PC1). Within each seed family, the herbicide resistant ‘R’ samples form separate clusters from their susceptible ‘S’ counterparts on PC2. The PC2 axis represents 12% of the total variance. In each seed family the ‘direction’ of separation of ‘R’ samples from ‘S’ is the same.

Differential expression analysis between ‘R’ and ‘S’ samples across the two seed families identified 643 differentially expressed genes. Of these, 341 were unique to family ‘CC2’, while 234 were unique to family ‘CC5’ (Figure 4b). A subset of 68 genes were found to be differentially expressed in both seed families. Hierarchical clustering of these 68 genes confirmed that resistance phenotype was a greater source of variability than seed family, and 81% (55) of these 68 genes were up-regulated in ‘R’ samples relative to ‘S’ for both families (Figure 4c).

**Figure 4.**
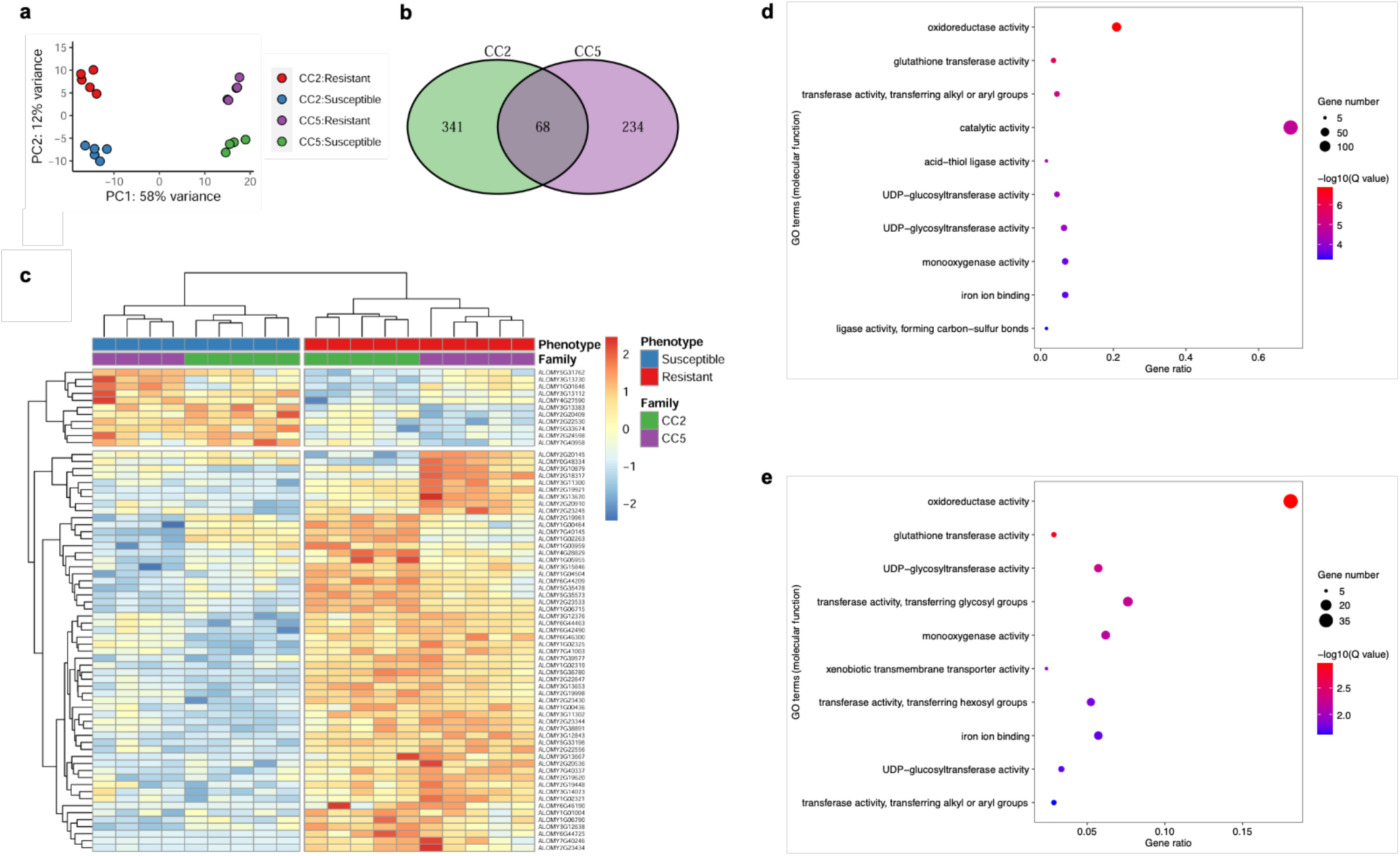
Differential gene expression analysis of the seed families CC2 and CC5, segregating for the NTSR herbicide resistance trait. **a**, Principal components analysis using all gene expression data. **b**, Numbers of differentially expressed genes comparing the ‘R’ and ‘S’ groups within each family. **c**, Heatmap and hierarchical clustering of the 68 differentially expressed genes consistently associated with NTSR across both seed families. **d** and **e**, Gene ontology terms, significantly overrepresented in the CC2 and CC5 families, respectively.

The list of 68 DEGs consistent across both seed families was found to contain three of eight previously recorded blackgrass NTSR candidate genes; ‘*AmGSTF1’*, ‘*AmGSTU2’*, and ‘*AmOPR1’* ^15,16^. In each case, expression of these three candidate genes was significantly higher in the ‘R’ phenotype than the ‘S’ (Supplementary Figure 6), agreeing with previously reported findings ^15,16^. Across the 68 consistent DEGs, six glutathione-S-transferases (GST), six cytochrome P450s, three ATP-binding cassette transporters (ABC transporters), and one aldo-keto reductase were found. This is consistent with a previous study which has demonstrated the potential importance of these key gene families in herbicide metabolism ^11^. Gene set enrichment analysis of DEGs for each family identified both shared and unique GO terms. Most of the shared overrepresented GO terms have been reported to be associated with NTSR, including glutathione transferase, UDP-glycosyltransferase, and some cytochrome P450 superfamilies. Xenobiotic transmembrane transporter was only overrepresented in CC5 (Figure 4d and 4e), indicating possible family-specific mechanism of resistance for CC5.

### Genetic coordination of NTSR via gene co-expression network analysis

Gene co-expression networks were constructed using traditional spearman-ranked and condition specific approaches that enable alternate strategies to examine the genetic coordination of NTSR mechanisms (Figure 5a and 5b, respectively). The traditional spearman ranked coefficient approaches resulted in a total of 16,601 nodes connected by 16,130 edges (Figure 5a). Hub gene sub-graphs display significant co-expressed gene interaction pairs that include candidate genes from the bulk segregant and RNA-seq studies. We identified a total of 13 CC2 specific sub-graphs and 20 for CC5 (Supplemental Figure 7a-d). In CC2, we found metabolism genes identified in the QTL-seq analysis, such as GST, aldo-keto reductase, and Beta-keto acyl synthase co-expressed with various transcription factors and other genes that could be involved in their regulation (Supplemental Figure 7a-b). An aldo-keto reductase was discovered through QTL-seq to be specific to the CC2 family that is also significantly upregulated in both the CC2 and CC5 families. The HMG transcriptional regulator is also positively correlated with two genes involved in metabolism: Tubulin/FtsZ family gene and a Ubiquitin carboxyl-terminal hydrolase, and negatively correlated with an Alpha-N-acetylglucosaminidase, (Supplemental Figure 7e). In the CC5 family sub-graphs, we identified alternate active genetic machinery that are co-expressed with genes identified in the QTL regions, such as Cytochrome p450s, thioesterase, glycosyl hydrolase, pectinesterase, exostensin gene family, and others connected with various classes of transporters and transcription factors/regulators (Supplemental Figure 7c-d). The condition specific network also partitioned clusters of co-expressed gene interactions pairs in both a family specific and overlapping manner. For example, this approach also identified an aldo-keto reductase and protein tyrosine/serine/threonine kinase unique to CC2. Oxioreductase, peroxidase, and vacuolar sorting were among CC5 specific clusters (Supplemental Figure 7f). This approach also identified a largely connected subgraph of connected genes discovered in both CC2 and CC5 bulk-segregant and RNA-seq analysis (Supplemental Figure 7g)

**Figure 5.**
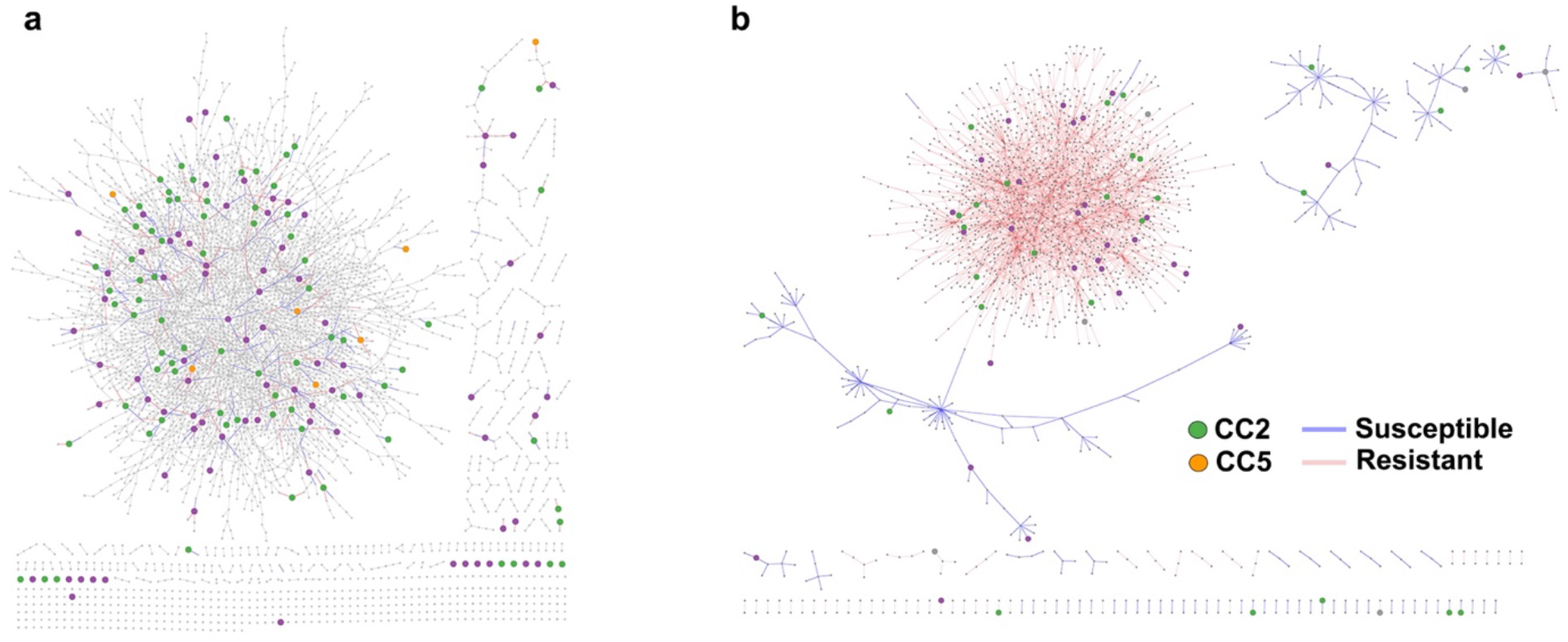
Genetic coordination of NTSR in CC2 and CC5. **a**, traditional spearman-ranked gene coexpression network derived from RNAseq expression that depicts common and unique genetic architecture underpinning NTSR in both the CC2 and CC5 families. Green nodes are unique to CC2, purple nodes are unique to CC5, and orange are common between both families. The graph was filtered for nodes with at least 2 connections. **b**, a condition-specific gene co-expression network derived from the RNAseq data taking into consideration plant phenotype (herbicide susceptible/resistant).

## Discussion

Despite the global distribution and impacts of weedy plants, few genomic resources have been developed to explore the genetics and evolution of weediness (see ^33^). Here, we present a reference-grade genome assembly for Europe’s worst agricultural weed ^21,23^. Metrics for genome size, completeness, structure, and quality indicate that this *A. myosuroides* genome is the largest and highest quality weedy plant genome produced to date. The very high proportion of sequence identified as transposable elements (TE), particularly LTR-retrotransposons, provides context for the genome size and plasticity that drives rapid evolution of weediness in blackgrass. It is well known that high levels of TE activity (transcription and movement) can provide the impetus for changes in gene expression, gene duplication, and genome organization, all of which can facilitate gene family evolution and the exaptation and co-option of genes to perform new functions, particularly in relation to biotic and abiotic stresses ^34^.

The analysis of genome structure and duplication identifies further signatures of a dynamically evolving and plastic genome as the basis for rapid adaptation in blackgrass. There is an over-representation of expanded gene families, evidence for a relatively recent unique genome duplication event in blackgrass, and a general excess of duplicated genes. It is notable that the paralogous genes associated with this duplication event are located on chromosomes 1, 2 and 3, where the densest regions of significant QTLs and differentially expressed genes are found (Fig 3). Also, expanded gene families included several gene functions that have been previously implicated in herbicide resistance and biotic and abiotic defense pathways. These features of the genome are consistent with a model that posits blackgrass weediness as an emergent property of a large and dynamic genome where rapid adaptation to new environmental stresses is enabled by exaptation of duplicated and differentially expressed genes under strong selective pressure ^35,36^.

Herbicides exert intense selection pressure on weed populations. The genomic basis of monogenic, target site resistance is well known ^11^, but with a few exceptions (for example ^37,38^), explorations of non-target site herbicide resistance (NTSR) have been limited to transcriptomics-based approaches, which do not inform the genetic basis of resistance directly. Using F_2_ seed families produced from two discrete NTSR genetic backgrounds, we demonstrate that NTSR is an oligogenic trait in blackgrass. Fifteen significant QTLs were identified (8 and 7 in the two seed families, respectively). Notably, there were no overlapping QTL regions between seed families derived from the two blackgrass populations, though significant QTLs were over-represented on chromosomes 2 and 3. These observations suggest that whilst selection for NTSR may be localized to certain regions of the genome, the genetic basis and genomic architecture of these traits is quite different amongst blackgrass populations. At the genomic level, evolution of NTSR amongst populations is divergent.

Our bulk segregant approach for sampling resistant and susceptible phenotypes identified a relatively small set of constitutively, differentially expressed genes (DEGs). As observed for the genomic data, there was a strong signal for selection of different molecular genetic mechanisms of resistance in the two resistant populations. 53% of DEGs were specific to one of the seed families, and 36% were specific to the other. However, there is also evidence for some convergence at the transcriptional level with 11% of the resistance associated DEGs being common to both seed families. It is difficult to definitively conclude from our analysis if the 89% (575) of family specific DEGs represent large functional differences in the NTSR phenotype, the effects of alternate paralogous gene expression, or perhaps pleiotropic differences arising from the different genetic background and genomic architecture of NTSR in the two parental NTSR populations. Our co-expression network analysis provides some indication, however, that whilst the metabolic machinery of NTSR across populations has common features, there are discrete genomic, transcriptional, and metabolic bases to NTSR in different blackgrass populations.

The common set of 68 DEGs included several genes and gene families that have been implicated in previous studies of NTSR in blackgrass ^13,14^ and in other weed species with evolved NTSR ^39-44^. The common DEGs are also over-represented on chromosomes 2 and 3 where most significant QTLs are found. We also note that a member of one of these gene families, the aldo-keto reductases, is closely linked to the most significant of our 15 QTL regions (chromosome 2, CC2 family). These differentially expressed gene families have roles in stress- and defense-related metabolism (Figures 4d and 4e) and are generally found to be expanded in blackgrass (Figures 2a and 2b). These findings add weight to our assertion that the rapid evolution of resistance and weediness in blackgrass is facilitated by its large, repetitive, and dynamic genome.

Access to a high-quality blackgrass genome has enabled us to identify several genomic features that can account for the weediness and adaptability of the species. Non-target site herbicide resistance is a complex trait that evolves repeatedly in blackgrass and other weedy plants, giving rise to a generalist resistance phenotype ^22^. Here, we clearly establish that it is an oligogenic trait, but that the genetic basis of NTSR can be markedly different between wild, evolved populations; albeit underpinned by some common metabolic pathways and manifested through genes that are similarly differentially expressed. Our results are consistent with those of Giacomini, et al. ^45^ who found physical clustering of differentially expressed genes, and whilst we do not find overlapping QTLs, there is strong evidence for selection of NTSR within similar genomic regions causing us to tentatively conclude that, as reported by Van Etten, et al. ^38^ and Kreiner, et al. ^37^, landscape scale evolution of NTSR likely results from both parallel and non-parallel patterns of evolution across the genome. These findings have wide significance for understanding the potential for rapid plant adaptation under novel and changing environments. They suggest that large and plastic plant genomes harbor sufficient standing genetic variation to enable rapid adaptation to novel stresses. The associated duplication and redundancy in plant genomes means that adaptation may not be mutation-limited and that the repeated evolution of resistance and/or tolerance relies on neither rare mutational events, nor hard selective sweeps.

## Methods

### Plant materials for genome sequencing and annotation

Blackgrass seeds collected in 2017 from section 8 of the Rothamsted ‘Broadbalk’ long-term experiment ^46^ were used to derive an individual blackgrass plant for genome sequencing. Established in 1843, these field plots have never received herbicide application, and extensive testing of this population (Rothamsted) over the last 20 years has confirmed that it remains susceptible to all herbicides, representing a true wild-type blackgrass genotype. In addition, two field-collected blackgrass seed populations (Peldon and Lola91) previously characterized as being strongly non-target-site resistance (NTSR) to acetyl-CoA carboxylase (ACCase) inhibiting herbicides were used to generate F_2_ seed families (named CC2 and CC5, respectively) for QTL-seq and RNA-seq analyses. Detailed protocols for the selection of a single herbicide sensitive plant for genome sequencing and for the development of CC2 and CC5 seed families is presented in the Supplementary Note.

### Genome survey

Previous study has reported that blackgrass has seven chromosomes ^27^. In this study, genome size was estimated through flow cytometry and k-mer based analysis. Flow cytometry was conducted on four field collected blackgrass populations (the Rothamsted, Lola91, and Peldon populations used within this study, along with a further herbicide susceptible population). Genome size estimates were generated for three replicate plants from each of these populations, against a known standard of the plant *Allium schoenoprasum*. Using these data, the blackgrass genome size was estimated as 3,312 – 3,423 Mb. K-mer based analysis from Illumina sequencing data derived from the Rothamsted population also indicated a genome size from 3,400 Mb to 3,550 Mb. We also estimated the heterozygosity and repeat content of the blackgrass genome with GCE package (https://github.com/BioInfoTools/GCE), the results suggest the blackgrass genome exhibits high level of heterozygosity (1.52%) and repeat content (84.2%). Due to the complexity of the blackgrass genome, we collected sequencing data from multiple sequencing platforms for genome assembling.

### Genome sequencing

#### Pacific Biosciences (PacBio) sequencing

high molecular weight (HMW) DNA was extracted from leaf tissues of a single plant (Rothamsted) that had been dark adapted for five days, used to construct PacBio SMRTbell libraries using SMRTbell Express Template Prep Kit 2.0, following the manufacturers’ protocols. SMRTbell libraries were sequenced on a PacBio Sequel II system and a total of six SMRT cells and 513 Gb (144 X coverage) data composed of ∼42 million reads were generated.

#### BioNano optical maps

HMW DNA was isolated from the same leaf tissue according to the BioNano Prep Plant Tissue DNA isolation protocol, and then fluorescently labelled using single-sequence-specific DLE1 endonucleases based on BioNano’s Direct Label and Stain (DLS) technology. The labelled DNA was loaded on the BioNano Genomics Saphyr system to scan by the sequencing provider. A total of 3,685,283 BioNano molecules were obtained with a total length of 860 Gb (241 X coverage).

#### Chromosome conformation capture sequencing by Hi-C

chromatin conformation capture data was generated using a Phase Genomics (Seattle, WA) Proximo Hi-C 2.0 Kit. Following the manufacturer’s instructions for the kit, intact cells from two samples were crosslinked using a formaldehyde solution, digested using the Sau3AI restriction enzyme, and proximity ligated with biotinylated nucleotides to create chimeric molecules composed of fragments from different regions of the genome that were physically proximal in vivo, but not necessarily genomically proximal. Continuing with the manufacturer’s protocol, molecules were pulled down with streptavidin beads and processed into an Illumina-compatible sequencing library. Sequencing was performed on an Illumina HiSeq 4000 system, yielding 126 Gb (35 X coverage) data.

#### Illumina short reads for polishing

DNA was extracted with the DNeasy Plant Mini Kit (QIAGEN) to prepare PCR-free paired-end libraries using the Illumina Genomic DNA Sample Preparation kit following the manufacturer’s instructions (Illumina). All paired-end libraries were sequenced on an Illumina NovaSeq 6000 system, generating 291 Gb (81 X coverage) of 150-nucleotide paired-end reads.

### Genome assembly

We performed de novo assembly of PacBio long reads into contigs with MECAT2 ^47^. This produced 12,107 contigs with an N50 of 0.9 Mb and a total size of 4,906 Mb. To improve the accuracy of the assembled contigs, two polishing strategies were performed including PacBio long reads polishing using Arrow program (https://github.com/PacificBiosciences/SMRT-Link) and Illumina short reads polishing using Pilon (v.1.20) ^48^. Polished contigs were repeat marked using WindowMasker ^49^ and then subjected to haplotype merging using HaploMerger2 ^50^ in terms of the heterozygosity of the blackgrass genome. BioNano data were first filtered based on molecule length (> 150Kb) and then aligned to primary contigs to select mapped molecules for de novo assembly to obtain the BioNano optical maps. The primary contigs and BioNano maps were combined to produce the hybrid scaffold assembly. The Hi-C reads were aligned to the hybrid scaffold assembly using the Juicer pipeline ^51^ and the hybrid scaffolds was then further scaffolded using the 3D-DNA pipeline ^52^. The results were manually examined using the Juicebox Assembly Tools, an assembly-specific module in the Juicebox visualization system ^53^. The Hi-C scaffolding resulted in seven pseudomolecule chromosomes. We performed gap filling using Cobbler (v0.6.1) ^54^ to eliminate the gaps generated in the scaffolding steps with PacBio long reads. In addition, the final assembled scaffolds were further polished using PacBio long reads with Arrow and Illumina short reads with Pilon ^48^. The detailed information is presented in the Supplementary Note.

### Genome assembly quality assessment

The quality of the assembled genome was evaluated by the following analyses. (1) The Illumina short reads used for polishing were mapped to the genome assembly using BWA-MEM, the mapping rate and genome coverage were examined. (2) The genome assembly was subjected to BUSCO (v.4.0.1) ^25^ analysis to assess the completeness of the assembly with the embryophyta_odb10 database. (3) The LRT Assembly Index ^26^ was calculated for assessing the genome assembly quality. (4) The assembled chromosome length was compared to the cytogenic chromosome length to check the correlation. The cytogenic chromosome length information has been reported in ^27^.

### Genome annotation

A comprehensive non-redundant repeat library for the blackgrass genome was built using EDTA, a de novo transposable element (TE) annotator that integrates structure- and homology-based approaches for TE identification ^55^. The EDTA pipeline incorporates LTRharvest, the parallel version of LTR_FINDER, LTR_retriever, GRF, TIR-Learner, HelitronScanner, and RepeatModeler as well as customized filtering scripts. Genome-wide prediction of ncRNAs, such as rRNA, small nuclear RNA and miRNA, was performed using INFERNAL software ^56^ with search against the Rfam database. The tRNA genes were predicted using tRNAscan-SE program ^57^.

Protein-coding genes were predicted by a combination of de novo prediction, homology-based and transcriptome-based strategies. SNAP ^58^, AUGUSTUS ^59^ and GeneMark ^60^ were used for ab initio gene predictions. For homology-based prediction, protein sequences of seven species (*A*.*thaliana, O*.*sativa, S*.*bicolor, B*.*distachyon, H*.*vulgare, Z*.*mays and T*.*aestivum*) were aligned to the genome assembly using GeMoMa program ^61^ to provide homology-based evidence. For transcriptome-based prediction, RNA-seq data were generated from the range of harvested blackgrass tissues (leaf, main stem, root, developing flowers, mature flowers pre-anthesis, and mature flowers with pollen). RNA-seq reads were processed to remove adapters and low-quality bases and assembled both de novo and genome guided using Trinity (v.2.4.0) ^62^ followed by the PASA program (http://pasapipeline.github.io) to improve the gene structures. All predicted gene structures were integrated into consensus gene models using EVidenceModeler ^63^. Functional annotation of protein-coding genes was carried out by comparing against SwissProt, GenBank nonredundant protein (NR), InterProScan and EggNOG databases. GO term for each gene was obtained from the corresponding InterPro descriptions. Additionally, the gene set was mapped to the KEGG pathway database using the online tool ‘BlastKOALA’ (https://www.kegg.jp/blastkoala/) ^64^.

### Long terminal repeat retrotransposons (LTR-RTs) insertion time estimation

As the direct repeat of an LTR-RT is identical upon insertion, the divergence between the LTR of an individual element reflects the time of the insertion. The insertion date (T) for each LTR-RT was computed by T = K/2μ, where K is the divergence rate and μ is the neutral mutation rate (K = - 3/4*ln(1-d*4/3), μ =1.3 × 10^−8^). Sequence identity (%) between the 5’ and 3’ direct repeats of an LTR candidate is approximated using blastn, so the proportion of sequence differences is calculated as d = 100% - identity%.

### Phylogeny and gene family

To identify orthologous and paralogous gene clusters, protein-coding genes from blackgrass and 11 other species (*A*.*tauschii, T*.*urartu, H*.*vulgare, P*.*tenuiflora, B*.*distachyon, O*.*sativa, Z*.*mays, S*.*bicolor, S*.*italica, E*.*haploclada, A*.*thaliana*) were analyzed using Orthofinder2 (v2.5.1) ^65^. In cases where there were multiple transcript variants, the longest transcript was selected to represent the coding region. A total of 476 single-copy orthologous genes were identified. Single-copy genes form each species were aligned using MUSCLE ^66^ and the alignments were concatenated. The concatenated alignment was subsequently used to construct a maximum likelihood phylogenetic tree using RAxML ^67^. The MCMCTree program ^68^ of PAML ^69^ was used to estimate the divergence time among 12 species. Three calibration time points were used based on previous publications and TimeTree website (http://www.timetree.org) as normal priors to restrain the age of the node, including 146-154 Mya between Arabidopsis and rice, 68-72 Mya between rice and sorghum, and 49-53 Mya between barley and Brachypodium. The gene family expansion and contraction were determined by comparing the gene cluster size differences between the ancestor and each species using CAFÉ program ^70^.

### Whole genome duplication and comparative genomic

To study the whole genome duplication events in the blackgrass genome, we performed the self-alignment within the blackgrass genome using LAST (v963) ^71^ and the syntenic blocks were identified using MCscanX ^72^. For each gene pair within syntenic blocks, the synonymous divergence levels (Ks) were calculated using the YN model in KaKs_Calculator ^73^. The Ks values of all gene pairs were plotted to identify the putative whole genome duplication events. To identify syntenic blocks between blackgrass and the other four species (*H*.*vulgare, A*.*tauschii, B*.*distachyon, O*.*sativa*), all-against-all BLASTP (E value < 1 × 10™5) was performed for the protein-coding gene set of each genome pair. Syntenic blocks were defined based on the presence of at least five synteny gene pairs using the MCScanX package ^72^ with default settings.

### QTL-seq (Bulk segregant analysis of SNPs)

Leaf tissue was harvested from the unsprayed tiller of all 25 ‘R’ and ‘S’ plants from each F_2_ family. In all cases, young leaf material was collected over one hour at midday, harvesting tissue from each plant into separate 5ml Eppendorf tubes. Each sample was immediately flash frozen in liquid nitrogen and stored at −80°C before use. For grinding, samples were kept cooled in liquid Nitrogen and homogenised using a micro-pestle. For bulk segregant analysis, four bulks were made by pooling DNA from all 25 selected individuals in each phenotypic group (herbicide resistant ‘R’, and susceptible ‘S’, in each of the CC2 and CC5 F_2_ families). Illumina paired-end reads were processed to remove adapters and low-quality sequences using Trimmomatic ^74^. Cleaned reads were mapped to the blackgrass reference genome using BWA. Variants were called using BCFtools (http://samtools.github.io/bcftools) and filtered using VCFtools (http://vcftools.sourceforge.net). QTL-seq pipeline was used for calculating the SNP-index, the ΔSNP-index was then calculated by subtracting the SNP-index of one bulk from that of another bulk ^30^.

### RNA-seq analysis

An RNA-seq analysis was also conducted using the 25-herbicide resistant ‘R’ and susceptible ‘S’ plants from identified from each F_2_ family. For each phenotypic group, five replicate RNA-bulks were created by pooling RNA from five individual plants. RNA was sequenced using standard Illumina TruSeq mRNAseq protocols. The quality of the RNA sequences derived from each sample was assessed using FastQC v0.11.8 ^75^ and preprocessed to remove the leading 10 bases from each read and any Illumina adapter sequences, together with any remaining reads shorter than 50 bases for adapters and low quality bases with Trimmomatic^74^. The trimmed reads for each sample were mapped to the *Alopecurus myosuroides* genome using Hisat2 v2.2.1 ^76^ with default parameters except for minimum alignment score parameters of L, 0, −0.6. Reads that mapped to coding sequences of annotated genes were counted using featureCounts v1.6.4 ^77^ with default settings. Differential gene expression between samples was analysed in R version 4.0.2 ^78^ using DeSeq2 ^79^.

The expression of all technical replicates was checked prior to analysis. First, all counts data were transformed using the regularised log-transform function ‘rlog()’ of the DESeq2 package. Transformed data were then visualised using both a principal components analysis (PCA), and hierarchical clustering of the Euclidean distance between samples. Visual inspection of these results identified one clear outlier sample (CC5 ‘S’ sample A), which was excluded from further analysis. A pre-filtering step was used to remove genes with zero or low counts before differential expression analysis. First, counts were summed across technical replicates to leave only biological samples. Next, genes were removed if they did not have at least one read per million in at least four samples (where four is equal to the minimum number of reps per treatment level) as per Anders, et al. ^80^. The filtered, biological replicates were analysed using the ‘DESeq()’ function of the DESeq2 package in R, specifying four phenotypic groups: CC2 ‘S’, CC2 ‘R’, CC5 ‘S’, and CC5 ‘R’. In total, 19,937 genes and 19 biological replicate samples were included in this analysis. To generate lists of differentially expressed genes (DEGs), specific comparisons were extracted for the ‘R’ vs ‘S’ samples within each family from this fitted model. Only genes which were significant (P<0.05) and with at least 1.5x fold difference in expression were categorised as differentially expressed. The resultant lists of DEGs for the CC2 and CC5 families were then intersected, to identify DEGs common to both.

Gene ontology information was combined from the Swissprot, Eggnog, and Interpro annotation files to create a single Gene:GO association map, containing 905,051 associations between 28,498 genes and 13,192 GO terms. Gene ontology enrichment analysis was performed for the DEGs using the ‘goseq()’ function of the goseq R package ^78^. The Gene:GO association map was specified as a custom gene category mapping to use for analysis, and enrichment scores for each gene ontology term were calculated using the Wallenius method (see Young, et al. ^81^). Resultant P-values were adjusted using the Benjamini and Hochberg method to further control the false discovery rate.

### Gene co-expression network construction

Trimmed means of M-values (TMM) were calculated from mapped RNAseq data using the edge-R package in R ^82^ to construct a gene expression matrix (GEM). The GEM was log_2_ transformed and quantile normalized using custom scripts in R ^78^. The traditional gene co-expression network (GCN) was created using the Knowledge Independent Network Construction tool (KINC v.3.4.0) ^83^. A gene correlation matrix was constructed using the Spearman rank correlation coefficient approach ^84^ with the following KINC specific parameters: --minsamp 15 –minexp -inf –mincorr 0.5 –maxcorr 0.99. A threshold for correlation was determined using the random matrix theory approach (RMT) in KINC with the following parameters: --tstart 0.95 –tstep 0.001 tstop 0.5 –threads 1 –epsilon 1e-6 –mineigens 50 –spline true –minspace 10 –maxpace 40 –bins 60 and was determined to be 0.919. The network was extracted using the extract function in KINC and visualized in Cytoscape v.3.9.0 ^85^. The condition specific GCN was constructed using the same GEM and Spearman ranked correlation coefficient approach in KINC, but also incorporated a Gaussian mixed model (GMM) to determine differentially expressed gene pair clusters that represent condition specific sub-graphs. Low powered edges were determined and filtered with the “corrpower” function in KINC with and alpha of 0.001 and power of 0.8. An annotation file was prepared in text format with samples either being annotated as “resistant” or “susceptible” and used to run the “cond-test”. Condition specific sub-graps were extracted and visualized in Cytoscape v.3.9.0 ^85^.

## Supporting information

Combined supplementary material

## Acknowledgements

DC, DM and PN were supported by the Smart Crop Protection Industrial Strategy Challenge Fund (grant no. BBS/OS/CP/000001) and Rothamsted Research as part of the Lawes Agricultural Trust. CS was supported by the Clemson University Research Fellows program. Rothamsted Research, Clemson University and Bayer Crop Science were equal contributors to costs associated with genomic and transcriptomic sequencing. The authors wish to thank Richard Hull and Laura Crook (Rothamsted Research, RR) for the growth and maintenance of plant material throughout this study, and David Hughes (RR) for bioinformatics assistance and advice.

## Author Contributions

PN, CS, and RB conceived the study and assembled project funding. CL, DC, and PN provided characterised plant material for sequencing. LC and CS assembled and annotated the blackgrass genome. LC, DC and CS analysed genomics and transcriptomics data sets and PN, DM and RB contributed to discussion and interpretation of data. LC, DC, CS and PN wrote the first draft of the paper and all authors contributed to subsequent editing and improvement.

## Data accessibility

** All sequence data will be archived in publicly available databases prior to final acceptance and publication of this article.

