## Supplementary material for "The blackgrass genome reveals patterns of divergent evolution of non-target site resistance to herbicides": Combined supplementary material

**Supplementary Note**

**The selection of sensitive individual plant for genome sequencing**

To identify an herbicide sensitive blackgrass individual for genome sequencing, 16 plants from the 'Rothamsted' population were grown from seed within a glasshouse until they reached approximately six vegetative tillers in size. Plants were physically split into five tillers (vegetative clones), and aboveground biomass trimmed to a uniform ~5cm length. Each tiller was re-potted and allowed to recover for one week in the glasshouse. A separate tiller from each plant was sprayed with the herbicidal actives; Fenoxaprop-p-ethyl (69 g ai $ha^{-1}$), Cycloxydim (150 g ai $ha^{-1}$), Sulfometuron (133 g ai $ha^{-1}$), and a commercial formulation of the two actives mesosulfuron-methyl and iodosulfuron-methyl (12 and 2.4 g ai $ha^{-1}$), while one further tiller from each plant remained unsprayed. Four weeks after spraying, all sprayed tillers were visually inspected to confirm sensitivity to the herbicide. Since all plants were confirmed as equally herbicide sensitive, a single unsprayed tiller was chosen at random from the screened plants to use as the source of tissue for genome sequencing.

The chosen plant was cultivated under glasshouse conditions until it had formed a large vegetative mass. At this point the plant was physically split once more into 24 tillers, which were again allowed to recover within the glasshouse until they had re-established as large vegetative plants. To ensure that only young, healthy tissue was harvested for sequencing, these well-established vegetative clones were cut back a final time to approximately 5cm of aboveground growth. After three weeks, the healthy aboveground re-growth from these plants was harvested and immediately flash frozen in liquid Nitrogen. By sampling in this manner, a large quantity (>50g) of freshly expanded, healthy leaf tissue was harvested from a single blackgrass plant to sequence as the basis of the blackgrass genome. All tissue was stored at -80°C before homogenization in bulk using liquid Nitrogen to keep tissue frozen.

To aid genome annotation, further individuals of the Rothamsted population were maintained within the glasshouse to maturity, with tissue collected across a range of phenological stages.

Healthy leaf tissue was harvested, along with 'stem' material from between the top two nodes of the main tiller. Flower heads were collected at three developmental stages; developing, un-emerged inflorescences, mature emerged flower heads pre-anthesis, and mature flower heads with anthers visibly extended and producing pollen. Blackgrass root tissue was also sampled from hydroponically grown plants. Briefly, pre-germinated seeds of the Rothamsted population were transferred to foam/mesh inserts and placed within 8 x 8 x 10cm magenta boxes (n=5 boxes, with six seeds per box), filled with 100ml of hydroponic media (Long-Ashton solution). Boxes were kept within an incubator (Sanyo MLR-350) at 17/11°C temperatures over a 14/10 hour light/dark cycle, and hydroponic solution was changed weekly. After four weeks, all root material was harvested and immediately flash frozen in liquid nitrogen and kept at -80°C before use.

**Genome assembling**

We adopted a hierarchical sequencing approach to generate a high-quality reference genome for blackgrass from a single individual using PacBio long reads, BioNano optical maps, chromatin conformation capture sequencing (Hi-C) and paired Illumina short reads (Supplementary Figure 1). First, we *de novo* assembled PacBio long reads into 12,107 contigs resulting in a total size of 4,906 Mb. After polishing, haplotype identification and deduplication, we obtained a primary contig set (representing homozygous regions of genome) that includes 7,866 contigs with an $N_{50}$ of 1.2 Mb. The total primary contig length is 3,475 Mb, which is consistent with our genome survey estimations based on flow cytometry and k-mer analysis (3,312-3,423 Mb and 3,400-3,550 Mb, respectively). BioNano data were first filtered based on molecule length (> 150Kb) and then aligned to primary contigs to select mapped molecules for de novo assembly, which produced 3,530 genome maps with an $N_{50}$ of 1.8 Mb for a total map length of 4367 Mb. Second, the primary contigs were corrected for false joins and scaffolded with optical map alignments (BioNano) to produce 2,512 hybrid scaffolds with the longest scaffold at 17.7 Mb. This increased the contiguity as indicated by $N_{50}$ value from 1.2 Mb to 2.3 Mb. Finally, hybrid scaffolds were ordered and oriented with Hi-C three-dimensional proximity data (Supplementary Figures 8 and 9). The final polished blackgrass genome assembly size was highly complete and contiguous at 3,572 Mb, including 3,400 (95.2%) Mb ordered as seven chromosomes with only 172 Mb of unanchored sequences.

**Generating $F_2$ families**

To explore the genetic basis of herbicide resistance in Blackgrass, two $F_2$ seed families were generated, with segregating non-target-site resistance (NTSR) to the Acetyl-CoA-Carboxylase inhibiting herbicide group (ACCase). To do this, plants from the herbicide susceptible Rothamsted population were grown in a glasshouse, along with individuals from two field-collected Blackgrass seed populations previously characterized as being strongly NTSR to the ACCase herbicides (Peldon, and Lola91). Plants were maintained for 2-3 months until well established. A single plant was chosen from each population and vegetatively cloned (tillering). The resistance phenotype was confirmed for these individuals by spraying one set of clones from each plant with a discriminatory dose of the herbicide Fenoxaprop-P-ethyl. The absence of ACCase target-site-resistance (TSR) in the selected plants was further confirmed by long-read sequencing of the ACCase gene.

To produce the $F_1$ generation, two clones of the herbicide sensitive individual from the Rothamsted population, and the resistant individual from each of the 'Peldon' and 'Lola91' populations, were vernalized for two months over winter in an unlit, unheated glasshouse. In the following spring, plants were paired such that each of the herbicide sensitive clones was paired with one of the two herbicide resistant individuals. Paired plants were moved to separate corners of a large glasshouse chamber, and before flowering, each pair was enclosed in a pollination bag. During flowering, bags were very gently shaken to assist pollen movement, facilitating cross-pollination of the two individuals enclosed. Once seed had matured, the pollination bags were removed, and seeds collected by gently shaking seed-heads into a paper envelope. As blackgrass is an obligate outcrossing species, seeds derived were the product of pairwise crossing between the enclosed pair of plants.

Collected seeds represent the $F_1$ generation of these two pairwise crosses. Seeds were air-dried at room temperature and cleaned using an air-column seed cleaner. Cleaned seeds were incubated at 30°C for three weeks to break dormancy. To generate the segregating $F_2$ families, 20 plants were grown from each of the $F_1$ lines. Plants were kept outside over Autumn – Spring allowing vernalization and vegetative establishment. Before flowering, the two sets of plants were moved into separate small pollination glasshouses. Plants were allowed to flower and openly cross-pollinate within each glasshouse, before collecting seeds in bulk from all individuals within each line. As before, these seeds were dried, cleaned and incubated at 30°C to break dormancy, and represent the $F_2$ segregating families used in the subsequent QTL-seq and RNA-seq analyses. Seed families were given the experimental codes 'CC2' ($F_2$ derived from Rothamsted x Peldon), and 'CC5' ($F_2$ derived from Rothamsted x Lola91). For an overview of $F_2$ line creation and subsequent bulk-segregant analysis, see Supplementary Figure 10.

**Material for bulked segregant analysis**

For each of the families (CC2 and CC5), 600 plants were grown under glasshouse conditions until well established, and split into separate vegetative tillers. For 300 plants in each family, two tillers were sprayed with a 'low' dose of the herbicide Fenoxaprop-p-ethyl. Plants which were consistently killed at this low dose represent herbicide susceptible individuals. Tillers for the remaining 300 plants of each family were sprayed with a 'high' doses of the herbicide. Consistent survivors of this high dose represent individuals with herbicide resistance. Herbicide doses were tailored to the two $F_2$ families based upon the resistance status of their original field-collected parental 'R' populations. For the CC2 family, doses of 69 g ai $ha^{-1}$ and 1104 g ai $ha^{-1}$ (1 and 16x the UK field rate) were found to optimally discriminate between the least- and most-resistant individuals, while for the CC5 family, doses of 138 g ai $ha^{-1}$ and 1656 g ai $ha^{-1}$ (2 and 24x field rate) were used. A single tiller from a further 24 randomly chosen plants in each line were sprayed at field rate with the herbicide cycloxydim, to provide an additional confirmation of the absence of TSR within these lines.

Plants were visually inspected on several occasions over the subsequent three weeks, and ranked to identify the 30-40 plants most affected by the 'low' doses (i.e. killed the quickest or at the smallest size), and least affected by the 'high' herbicide doses (i.e. those with vigorous growth matching the unsprayed tiller). At each dose, both tillers from the 30-40 chosen individuals were harvested, alongside both tillers from a further 100 randomly chosen plants. Aboveground 'fresh' biomass was recorded for each tiller, with dry biomass recorded following oven drying at 80°C for 48 hours. Biomass measurements were used in combination with the visual scoring information to select the 25 plants either most susceptible ('S'), or most resistant ('R') to the herbicide within each family. Leaf material from the un-sprayed clones of these chosen 'R' and 'S' plants was harvested, flash frozen in liquid Nitrogen, and used throughout the RNA-seq and QTL-seq bulked segregant analyses (Supplementary figure 10).

**Confirming resistance status of the $F_2$ families**

A glasshouse phenotyping assay was used to confirm the resistance status of the $F_2$ seed lines in comparison to their original parental populations. Seeds of both the 'CC2' and 'CC5' $F_2$ families, as well as the original 'Rothamsted', 'Peldon', and 'Lola91' populations, were germinated in Petri dishes containing three Whatman No. 1 filter papers soaked in 5 ml of 0.02 M KNO3. Petri dishes were incubated for seven days in a Sanyo MLR-350 growth cabinet, with a 17/11°C temperature cycle and a 14/10 hour light/dark cycle. Germinated seeds were transplanted 5 – 10mm below the soil surface in 8 cm plastic plant pots (six seeds per pot), filled with a Kettering loam soil mixed with 2 kg m-2 osmocote fertiliser. Plants were maintained in a glasshouse at approximately 16/10°C until seedlings had reached the three leaf stage, before spraying with the ACCase herbicide Fenoxaprop-P-ethyl (using the commercial formulation 'Polecat'). Doses used were: 0, 2.16, 4.31, 8.63, 17.25, 34.5, 69, 138, 276, 552, and 1104 g ai $ha^{-1}$, representing 0, 0.03, 0.06, 0.125, 0.25, 0.5, 1, 2, 4, 8, and 16x the UK field rate for this active. Herbicide was applied using a fixed track sprayer, with a spray nozzle (flat fan 110015VK; Teejet, Wisbech, UK) mounted 50 cm above the pots, and boom speed set at 0.33 m $s^{-1}$, applying herbicide at a rate of 199 l $ha^{-1}$. Three pots of each population (n=3) were sprayed per dose, and pots were placed back in the glasshouse immediately following spraying. Three weeks after herbicide application, all plants were assessed for mortality, and above-ground leaf and shoot biomass was harvested, dried at 80˚C for 48 hours, and weighed. Dose-response relationships were compared for each $F_2$ line relative to their parent populations. In both cases, the phenotypic level of herbicide resistance in the two $F_2$ families was intermediate, between that of the resistant and susceptible parental families, as expected for inheritance of this potentially quantitative trait (Supplementary Figure 11 a, b).

**Confirming the ‘R’ and ‘S’ phenotypes in the $F_3$ generation**

After tissue sampling, the 25 confirmed ‘R’ and ‘S’ plants from the CC2 and CC5 $F_2$ families were re-potted into larger, 6-inch pots, and grown to maturity. At the onset of flowering, plants were moved into four small pollination glasshouses, with all 25 plants from the same phenotypic group (CC2 ‘R’, CC2 ‘S’, CC5 ‘R’, CC5 ‘S’) kept together. Flowering heads were agitated once per week to help pollen movement and facilitate bulk cross pollination within each glasshouse. Seeds were collected at maturity and air dried at room temperature before cleaning with an air column seed cleaner to remove empty husks and debris. Cleaned seeds were incubated at 30°C in a Sanyo MLR-350 growth cabinet for three weeks to break dormancy.

These seeds represent $F_3$ lines from the CC2 and CC5 families. If the ‘R’ and ‘S’ plants identified from the $F_2$ truly represent individuals with genetic segregation for the herbicide resistance trait, we would expect that a difference in ‘R’ and ‘S’ phenotype would be maintained into this $F_3$ generation. To confirm this, seeds from these ‘R’ and ‘S’ $F_3$ lines, along with the $F_2$ CC2 and CC5 lines that they were derived from, were tested under glasshouse conditions for their Fenoxaprop-P-ethyl dose-response relationship. Seeds were germinated in Petri dishes containing three Whatman No. 1 filter papers soaked in 5 ml of 0.02 M KNO3. Petri dishes were incubated for seven days in a Sanyo MLR-350 growth cabinet, with a 17/11°C temperature cycle and a 14/10 hour light/dark cycle. Six germinated seeds were transplanted 5 – 10mm below the soil surface into an 8 cm plastic plant pot containing a Kettering loam soil mixed with 2 kg m-2 osmocote fertilizer.

Pots were kept in a glasshouse for approximately three weeks until plants had reached the three-leaf stage, and were then sprayed with Fenoxaprop-P-ethyl using the same commercial formulation as previously (Foxtrot). Doses used were 0, 1.08, 2.16, 4.31, 8.63, 17.25, 34.5, 69, 138, 276, 552, 1104, and 2208 g ai $ha^{-1}$, and for each line, three replicate pots were sprayed per dose. The same fixed track-spraying unit and Teejet 110015VK nozzle were used, as previously described. Three weeks after spraying, plants were visually assessed for survival and dose-response relationships compared. Comparison of herbicide survival confirms that the expected 'R' and 'S' phenotype is maintained within these 'R' and 'S' $F_3$ lines (Supplementary Figure 11, c, d).

Supplementary Table 1. Genome size estimation based on flow cytometry.

| Sample ID | Population | Phenotype | DNA content (pg/2C) | Mbp/2C | Gbp/1C |
|---|---|---|---|---|---|
| BG1 | Rothamsted | S | 6.90 | 6750 | 3.37 |
| BG2 | Rothamsted | S | 6.74 | 6595 | 3.30 |
| BG3 | Rothamsted | S | 6.81 | 6656 | 3.33 |
| BG4 | PELD | R | 6.77 | 6625 | 3.31 |
| BG5 | PELD | R | 7.54 | 7373 | 3.69 |
| BG6 | PELD | R | 6.77 | 6625 | 3.31 |
| BG7 | LOLA 81 | S | 6.84 | 6687 | 3.34 |
| BG8 | LOLA 81 | S | 6.74 | 6595 | 3.30 |
| BG9 | LOLA 81 | S | 7.00 | 6846 | 3.42 |
| BG10 | LOLA 91 | R | 6.90 | 6750 | 3.37 |
| BG11 | LOLA 91 | R | 6.77 | 6625 | 3.31 |
| BG12 | LOLA 91 | R | 6.87 | 6718 | 3.36 |
| Average | NA | NA | 6.89 | 6737 | 3.37 |

Supplementary Table 2. Genome size estimation based on k-mer analysis.

| ***k*-mer length** | **Total number of *k*-mer** | ***k*-mer coverage** | **Estimated genome size** |
|---|---|---|---|
| 17 | 237,986,849,712 | 70 | 3400Mb |
| 19 | 234,416,794,205 | 68 | 3447Mb |
| 21 | 230,846,540,235 | 66 | 3498Mb |
| 31 | 212,995,740,384 | 60 | 3550Mb |

Supplementary Table 3. Statistics of the assembled seven chromosomes of *A. myosuroides*.

| Chromosome | Anchored scaffold number | Length of chromosome |
|---|---|---|
| Chr1 | 627 | 781,714,917 |
| Chr2 | 416 | 571,710,514 |
| Chr3 | 350 | 522,029,352 |
| Chr4 | 321 | 443,909,942 |
| Chr5 | 278 | 369,436,349 |
| Chr6 | 264 | 359,679,614 |
| Chr7 | 238 | 351,570,514 |
| Total | 2494 | 3,400,051,202 |

Supplementary Table 4. BUSCO analysis of genome completeness.

| Description | Gene number | Percentage (%) |
|---|---|---|
| Complete BUSCOs (C) | 1563 | 96.9 |
| Complete and single-copy BUSCOs (S) | 1497 | 92.8 |
| Complete and duplicated BUSCOs (D) | 66 | 4.1 |
| Fragmented BUSCOs (F) | 14 | 0.9 |
| Missing BUSCOs (M) | 37 | 2.2 |
| Total BUSCO groups searched | 1614 | 100 |

Supplementary Table 5. Statistics of the annotated transposon elements (TEs).

| Class | Superfamily | Number | Total size (bp) | Percentage (%) |
|---|---|---|---|---|
| Class 1 | LTR-retrotransposons | | | |
| | *Gypsy* (RLG) | 1922808 | 1,367,269,327 | 39.16 |
| | *Copia* (RLC) | 445775 | 298,678,755 | 8.56 |
| | Unclassified (RLX) | 1574609 | 623,678,411 | 17.86 |
| | Non-LTR retrotransposons | | | |
| | Long interspersed nuclear elements (RIX) | 20527 | 12,053,133 | 0.35 |
| | Short interspersed nuclear elements (SIX) | 2807 | 797,889 | 0.02 |
| Class 2 | DNA transposons | | | |
| | hAT (DTA) | 41471 | 15,852,349 | 0.45 |
| | CACTA (DTC) | 335408 | 169,435,014 | 4.85 |
| | Harbinger (DTH) | 124608 | 61,410,666 | 1.76 |
| | Mutator (DTM) | 181317 | 79,309,130 | 2.27 |
| | Mariner (DTT) | 189514 | 60,919,894 | 1.74 |
| | Unclassified (DXX) | 24210 | 11,107,295 | 0.32 |
| | Helitron (DHH) | 248002 | 109,086,060 | 3.12 |
| Others | XXX | 176175 | 41,788,046 | 1.20 |
| Total | TEs | 5287231 | 2,851,385,969 | 81.68 |

Supplementary Table 6. Summary of identified QTLs from CC2 and CC5 populations.

| Population | Chromosome | Name | Start(bp) | End(bp) | Length(bp) | Peak delta SNP index | Peak position (bp) | Average delta SNP index |
|---|---|---|---|---|---|---|---|---|
| CC2 | Chr2 | *qtl-cc2-2-1* | 78,155,610 | 80,414,345 | 2,258,735 | 0.35 | 79,286,939 | 0.31 |
| | Chr2 | *qtl-cc2-2-2* | 84,968,723 | 86,789,597 | 1,820,874 | 0.33 | 85,985,573 | 0.31 |
| | Chr2 | *qtl-cc2-2-3* | 155,616,708 | 158,148,961 | 2,532,253 | 0.41 | 156,325,090 | 0.39 |
| | Chr3 | *qtl-cc2-3-1* | 314,531,189 | 315,950,139 | 1,418,950 | 0.30 | 315,049,343 | 0.27 |
| | Chr5 | *qtl-cc2-5-1* | 190,968,113 | 192,166,507 | 1,198,394 | 0.29 | 191,929,219 | 0.27 |
| | Chr5 | *qtl-cc2-5-2* | 210,023,549 | 211,392,868 | 1,369,319 | -0.29 | 210,693,320 | -0.26 |
| | Chr6 | *qtl-cc2-6-1* | 139,536,063 | 142,546,311 | 3,010,248 | 0.28 | 140,494,464 | 0.27 |
| CC5 | Chr2 | *qtl-cc5-2-1* | 28,124,368 | 30,262,000 | 2,137,632 | -0.33 | 29,036,282 | -0.30 |
| | Chr3 | *qtl-cc5-3-1* | 197,701,391 | 198,108,389 | 406,998 | -0.30 | 197,792,990 | -0.27 |
| | Chr3 | *qtl-cc5-3-2* | 224,118,449 | 226,070,109 | 1,951,660 | 0.30 | 224,317,480 | 0.29 |
| | Chr3 | *qtl-cc5-3-3* | 229,745,161 | 230,705,090 | 959,929 | 0.29 | 230,421,617 | 0.28 |
| | Chr3 | *qtl-cc5-3-4* | 248,724,538 | 251,387,988 | 2,663,450 | -0.35 | 249,793,371 | -0.33 |
| | Chr3 | *qtl-cc5-3-5* | 281,782,841 | 288,136,846 | 6,354,005 | -0.39 | 283,456,062 | -0.35 |
| | Chr3 | *qtl-cc5-3-6* | 308,719,007 | 313,553,107 | 4,834,100 | -0.41 | 309,948,215 | -0.37 |
| | Chr3 | *qtl-cc5-3-7* | 444,582,369 | 447,919,982 | 3,337,613 | -0.41 | 446,569,526 | -0.37 |

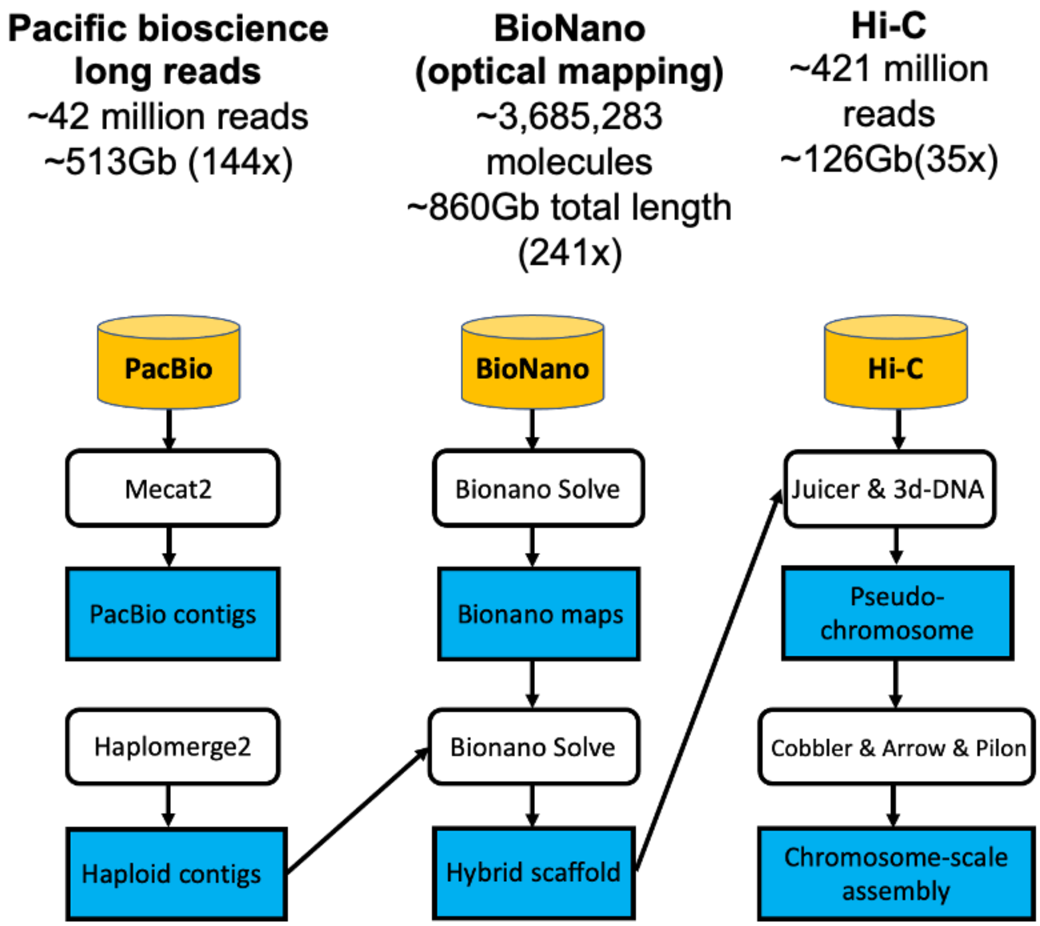


Supplementary Figure 1. Pipeline of genome assembly for the blackgrass.

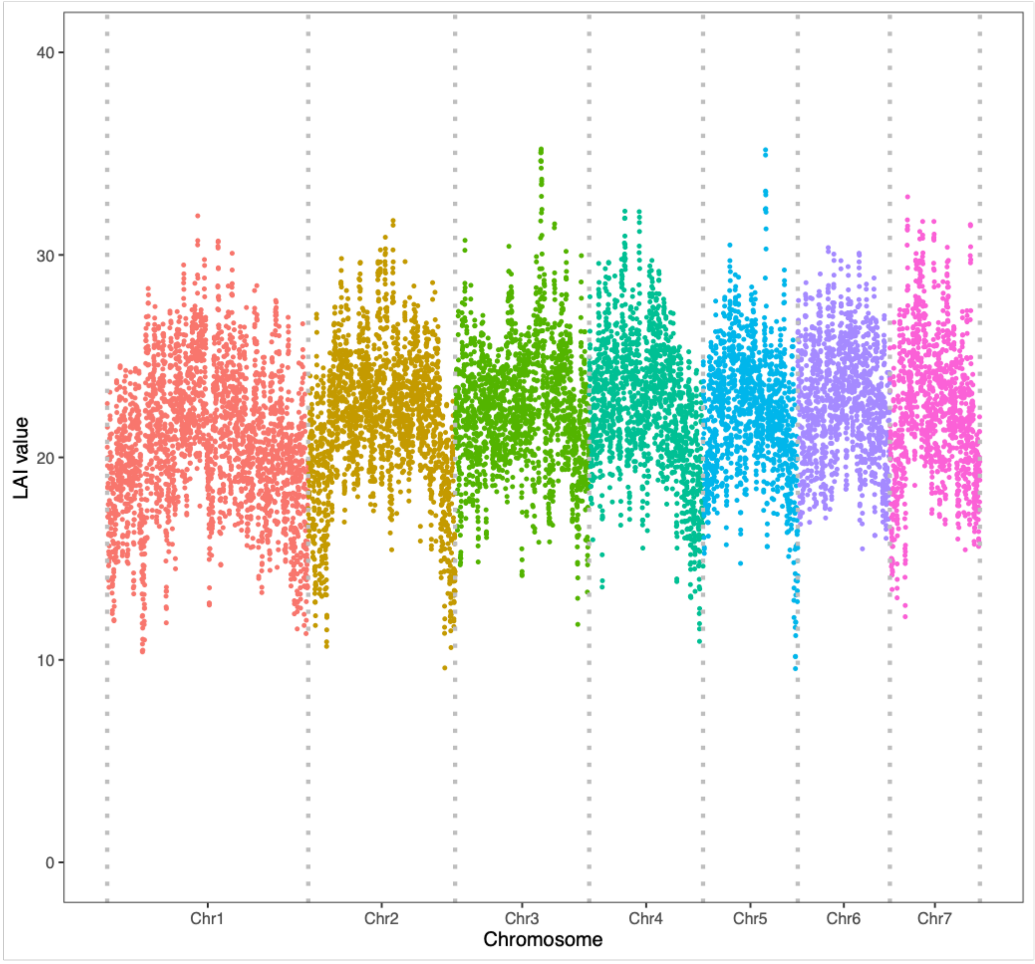


Supplementary Figure 2. Long terminal repeat assembly index (LAI) values across seven assembled blackgrass chromosomes.

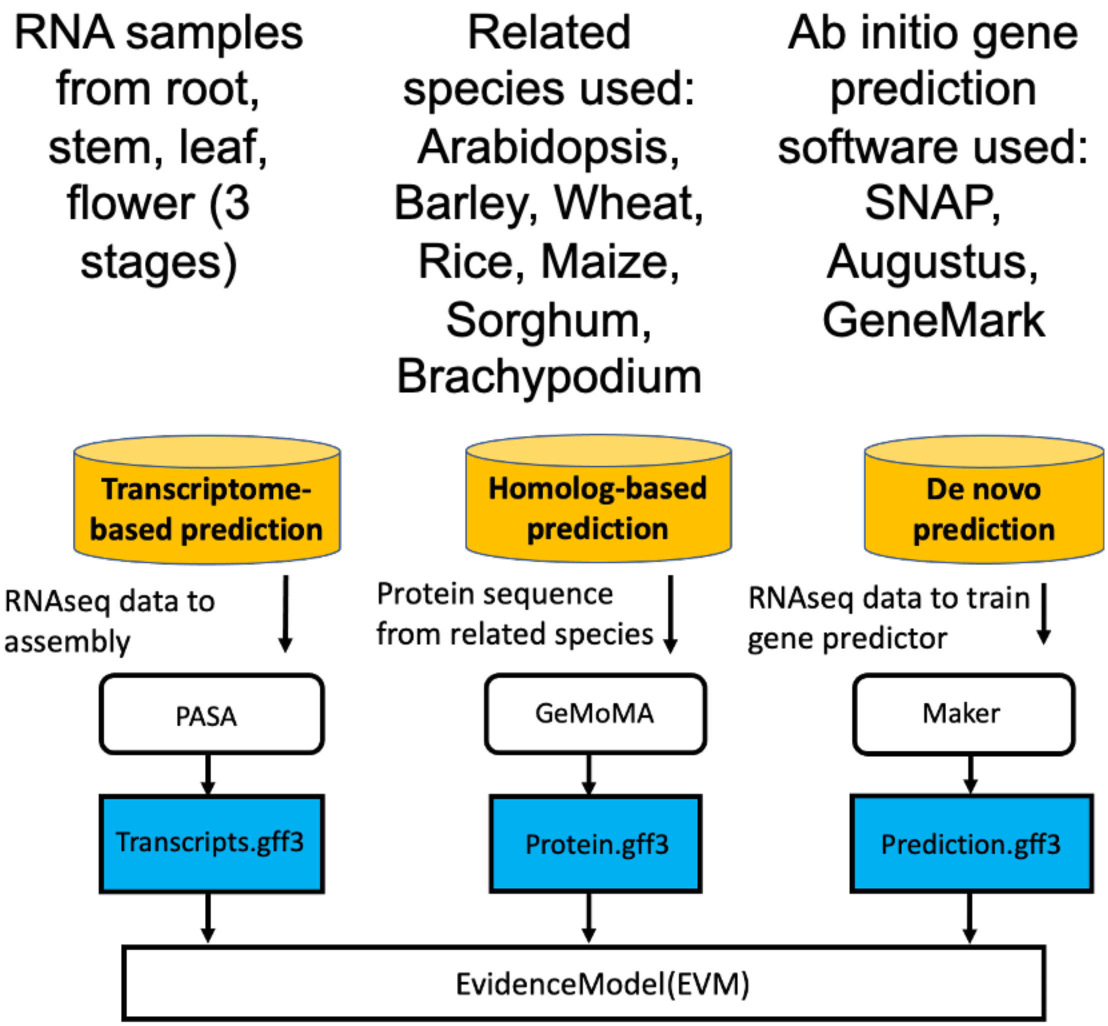


Supplementary Figure 3. Pipeline of gene annotation for the blackgrass.

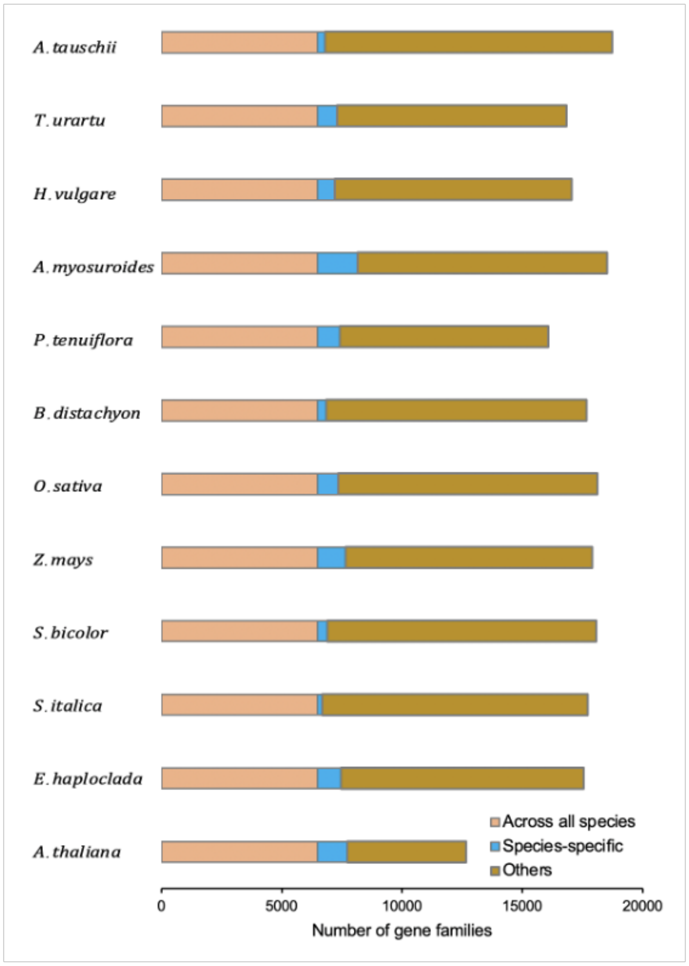


Supplementary Figure 4. The distribution of different types of gene family, including the ones presented across all the species, species-specific and all other types.

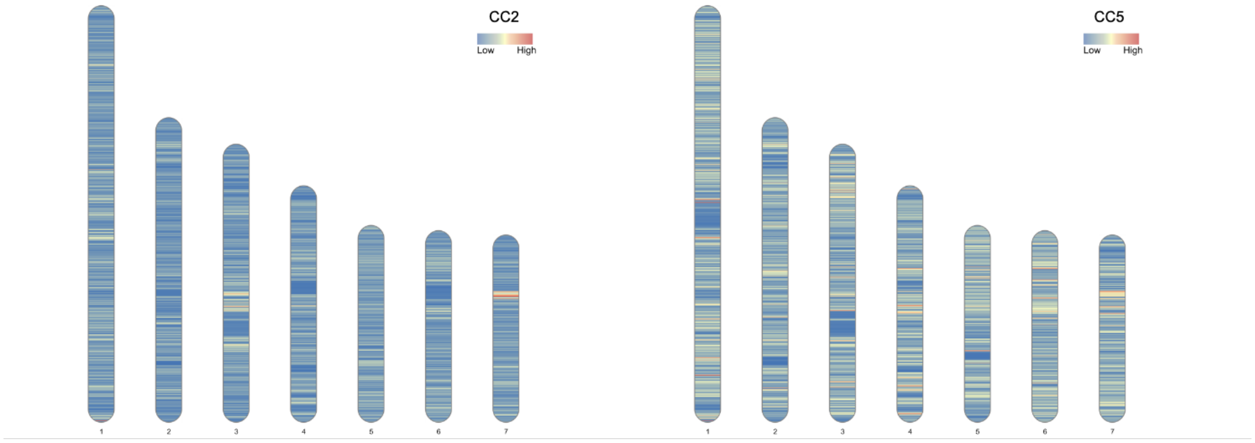


Supplementary Figure 5. SNP marker density for CC2 and CC5 population, respectively.

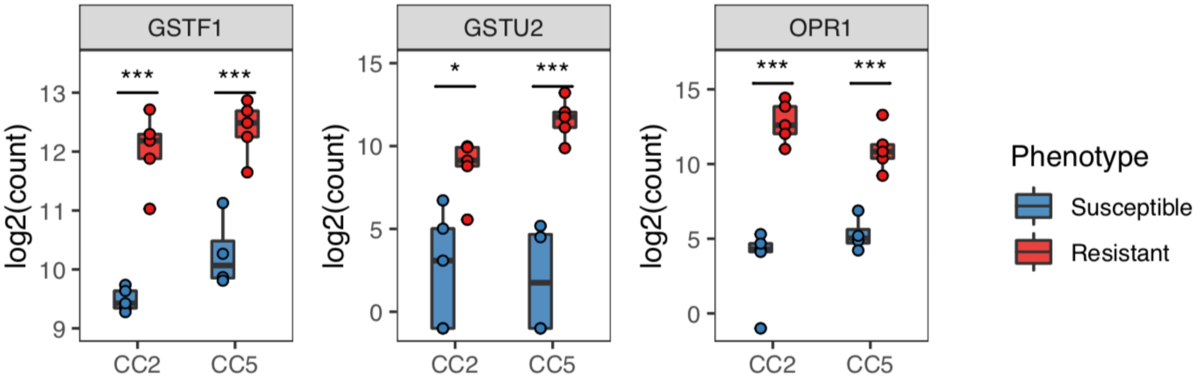


Supplementary Figure 6. Differential expression of three previously reported NTSR candidate genes.

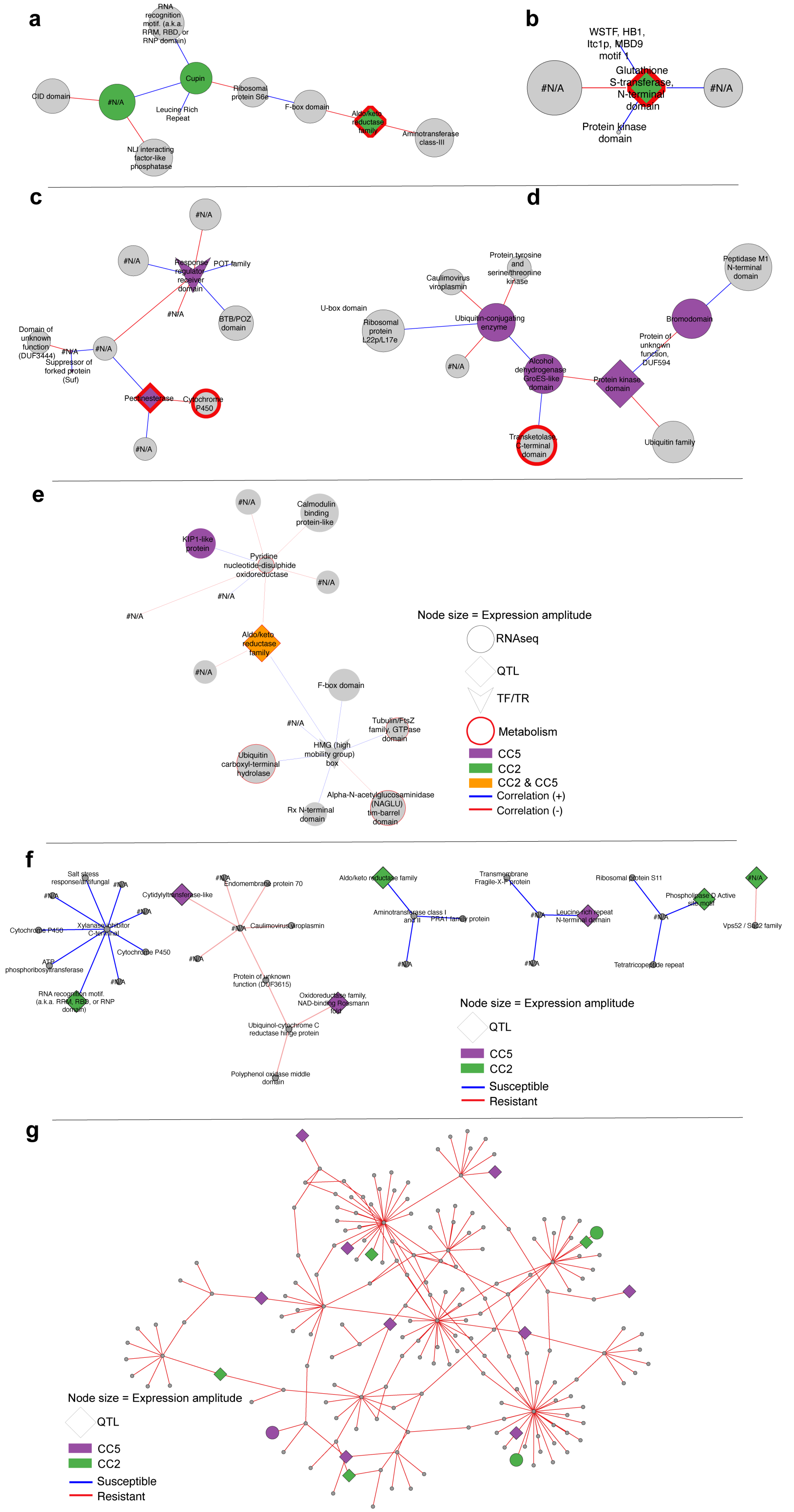
a
RNA recognition motif. (a.k.a. RRM, RBD, or RNP domain)
Cupin
CID domain
#N/A
Leucine Rich Repeat
Ribosomal protein S6e
F-box domain
Aldo/keto reductase family
Aminotransferase class-III
NLI interacting factor-like phosphatase
b
WSTF, HB1, Itc1p, MBD9 motif 1
Glutathione S-transferase, N-terminal domain
#N/A
#N/A
Protein kinase domain
c
#N/A
#N/A
Response regulator receiver domain
POT family
#N/A
BTB/POZ domain
Domain of unknown function (DUF3444)
#N/A
#N/A
Suppressor of forked protein (Suf)
Pectinesterase
Cytochrome P450
#N/A
d
Protein tyrosine and serine/threonine kinase
Caulimovirus viroplasmin
Peptidase M1 N-terminal domain
U-box domain
Ribosomal protein L22p/L17e
Ubiquitin-conjugating enzyme
Bromodomain
Protein of unknown function, DUF594
#N/A
Alcohol dehydrogenase GroES-like domain
Protein kinase domain
Transketolase, C-terminal domain
Ubiquitin family
e
#N/A
Calmodulin binding protein-like
KIP1-like protein
Pyridine nucleotide-disulphide oxidoreductase
#N/A
#N/A
#N/A
Aldo/keto reductase family
#N/A
F-box domain
#N/A
Tubulin/FtsZ family, GTPase domain
HMG (high mobility group) box
Ubiquitin carboxyl-terminal hydrolase
Alpha-N-acetylglucosaminidase (NAGLU) tim-barrel domain
Rx N-terminal domain
Node size = Expression amplitude
RNAseq
QTL
TF/TR
Metabolism
CC5
CC2
CC2 & CC5
Correlation (+)
Correlation (-)
f
Salt stress response/antifungal
#N/A
#N/A
#N/A
Xylanase inhibitor C-terminal
Cytochrome P450
Cytochrome P450
ATP phosphoribosyltransferase
#N/A
RNA recognition motif. (a.k.a. RRM, RBD, or RNP domain)
Cytidylyltransferase-like
#N/A
Endomembrane protein 70
#N/A
Caulimovirus viroplasmin
#N/A
Protein of unknown function (DUF3615)
Oxidoreductase family, NAD-binding Rossmann fold
Ubiquinol-cytochrome C reductase hinge protein
Polyphenol oxidase middle domain
Aldo/keto reductase family
Aminotransferase class I and II
PRA1 family protein
#N/A
Transmembrane Fragile-X-F protein
#N/A
Leucine rich repeat N-terminal domain
#N/A
Ribosomal protein S11
Phospholipase D Active site motif
#N/A
#N/A
Tetratricopeptide repeat
#N/A
Vps52 / Sac2 family
Node size = Expression amplitude
QTL
CC5
CC2
Susceptible
Resistant
g
Node size = Expression amplitude
QTL
CC5
CC2
Susceptible
Resistant

**Supplementary Figure 7. Parallel and overlapping sub-graphs in CC2 and CC5. A-b,** Hub genes unique to CC2 subset from the traditional network discovered in the bulk-segregant analysis. **c-d,** Hub genes unique to CC5 subset from the CC5 traditional network, and **e,** a hub gene that was discovered in both the CC2 bulk segregant analysis and is differentially expressed in resistant individuals in both families. **F.** Representative condition-specific sub-graphs that display both independent and common gene-interaction pairs between CC2 and CC5 families. **G.** A representative sub-graph from the condition specific network analysis that shows a large subgraph of genes identified in the bulk segregant analysis.

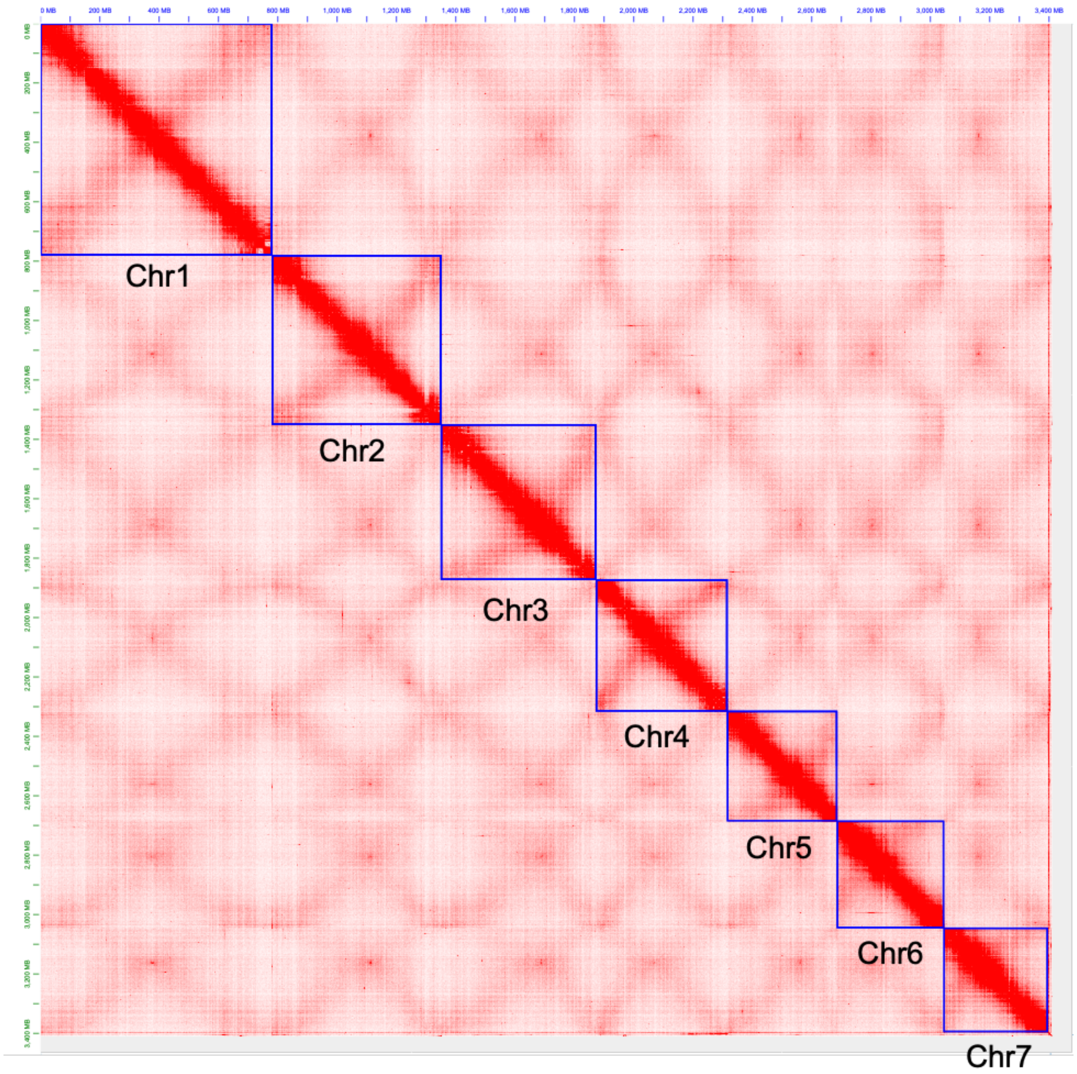


Supplementary Figure 8. Snapshot of Juicebox Assembly Tools output after manual correction.

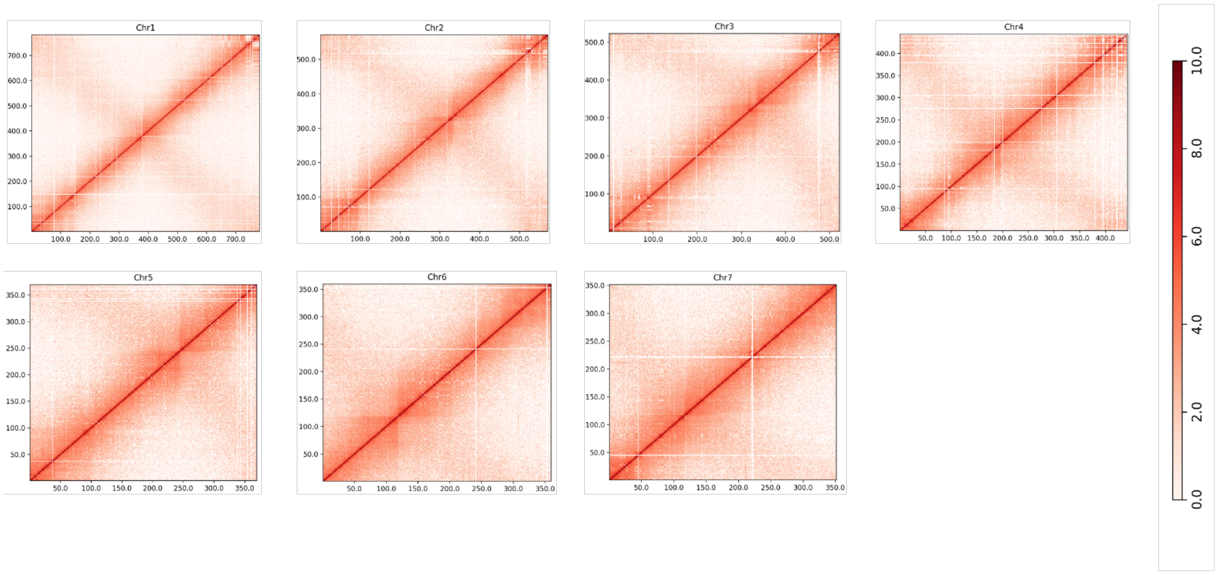


Supplementary Figure 9. Genome-wide analysis of chromatin interactions at 1-Mb resolution in the assembled blackgrass genome.

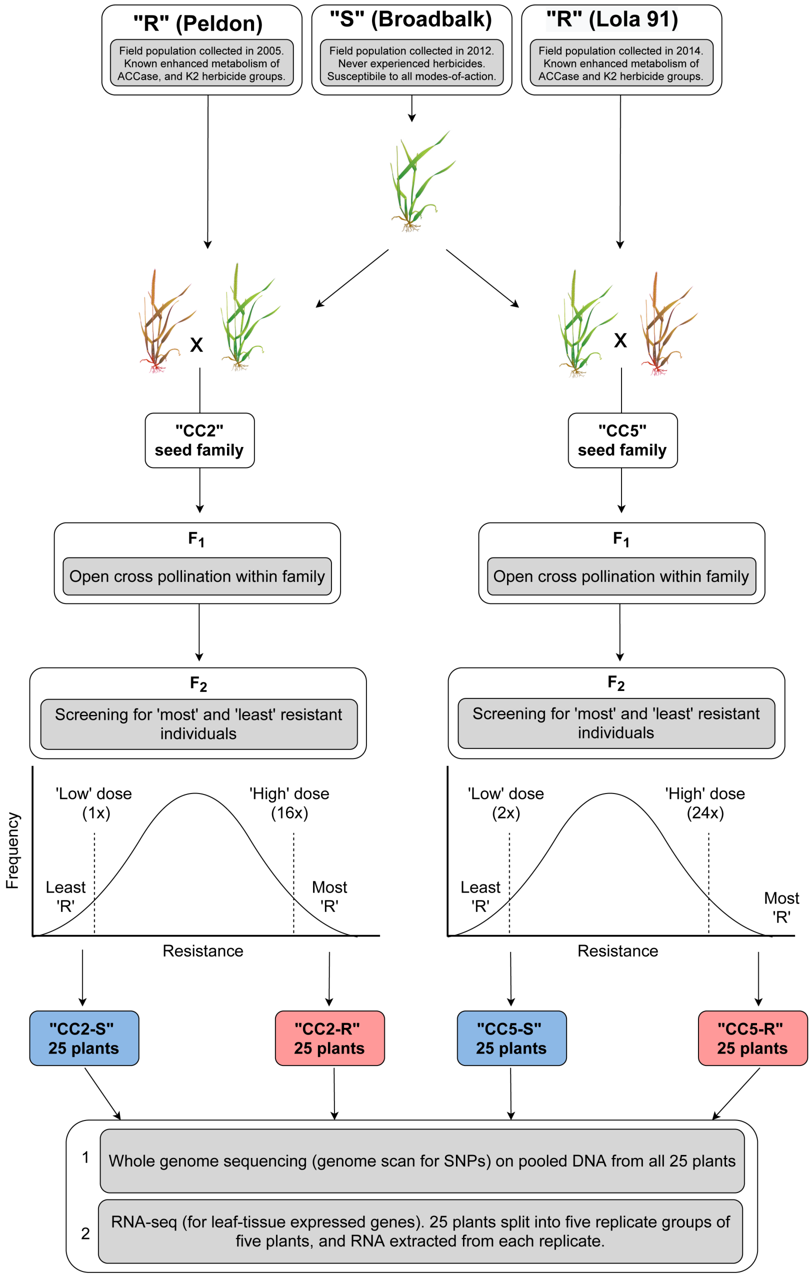


Supplementary Figure 10. Overview of the steps involved in creating the experimental seed families. Initially a single ‘S’ plant of the Broadbalk population was vegetatively cloned and cross pollinated with an individual from one of the two ‘R’ populations. Herbicide screening in the $F_2$ generation was performed using the ACCase inhibitor Fenoxaprop-P-ethyl. Tissue was collected from 25 plants identified as either ‘R’ or ‘S’ in each family, and used for whole genome sequencing and analysis of gene expression.

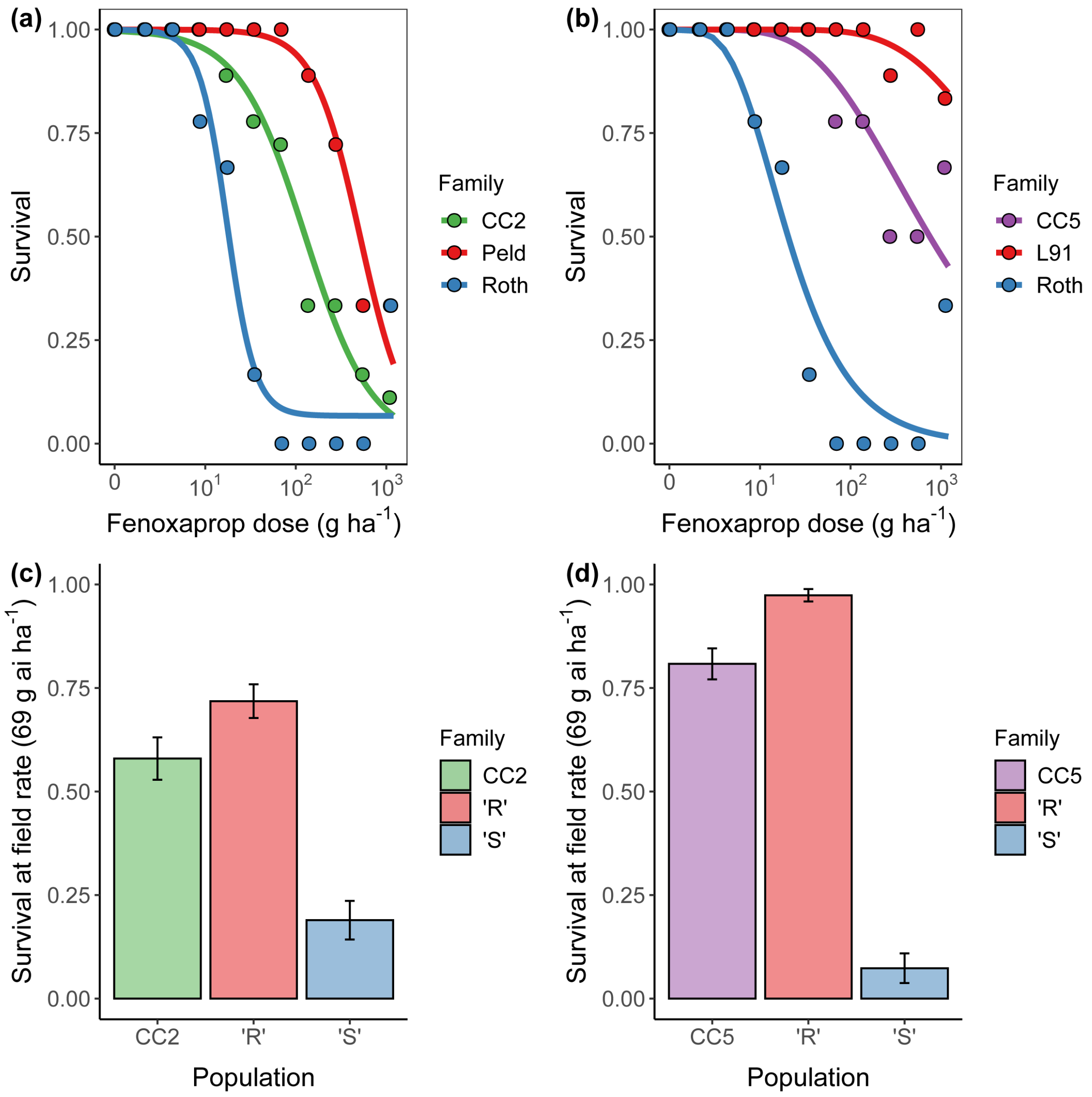


Supplementary Figure 11. Confirmation of resistance phenotype in the experimental seed families. **a** and **b** show the dose-response relationship of the two $F_2$ seed families (CC2 and CC5), relative to their respective parental populations. **c** and **d** show herbicide resistance of progeny from the ‘R’ and ‘S’ bulks, relative to the seed family from which they were derived. In all cases resistance was determined using a commercial formulation of the Acetyl CoA carboxylase inhibiting herbicide fenoxyprop-P-ethyl.
